# Optimizing Phosphopeptide Structures That Target 14-3-3ε in Cutaneous Squamous Cell Carcinoma

**DOI:** 10.1101/2023.10.03.560749

**Authors:** Seraphine Kamayirese, Sibaprasad Maity, Lynne M. Dieckman, Laura A. Hansen, Sándor Lovas

## Abstract

14-3-3ε is involved in various types of malignancies by increasing cell proliferation, promoting cell invasion or inhibiting apoptosis. In cutaneous squamous cell carcinoma (cSCC), 14-3-3ε is over expressed and mislocalized from the nucleus to the cytoplasm where it interacts with the cell division cycle 25 A (CDC25A) and suppresses apoptosis. Hence inhibition of the 14-3-3ε - CDC25A interaction is an attractive target for promoting apoptosis in cSCC. In this work, we optimized the structure of our previously designed inhibitor of 14-3-3ε – CDC25A interaction, pT, a phosphopeptide fragment corresponding to one of the two binding regions of CDC25A to 14-3-3ε. Starting from pT, we developed peptide analogs that bind 14-3-3ε with nanomolar affinities. Peptide analogs were designed by shortening the pT peptide, and introducing modifications at position 510 of the pT(502-510) analog. Both molecular dynamics (MD) simulations and biophysical methods were used to determine peptides binding to 14-3-3ε. Shortening the pT peptide from 14 to 9 amino acid residues resulted in a peptide (pT(502-510)) that binds 14-3-3ε with a K_D_ value of 45.2 nM. Gly to Phe substitution in position 510 of pT(502-510) led to further improvement in affinity (K_D_: 22.0 nM) of the peptide for 14-3-3ε. Our results suggest that the designed peptide analogs are potential candidates for inhibiting 14-3-3ε -CDC25A interactions in cSCC cells; thus, inducing their apoptosis.

## 1. INTRODUCTION

14-3-3 proteins are a family of highly conserved proteins that are expressed in all eukaryotic cells. In mammals there are seven isoforms, β (when phosphorylated, β is called α) γ, ε, ζ (when phosphorylated, ζ is called δ), σ, η and τ ^1–4^. 14-3-3 proteins are biologically active in either homo- or heterodimeric forms, and the 14-3-3ε isoform only exists in heterodimeric form^5^. Each monomer of 14-3-3 consists of nine antiparallel α-helices (αA, αB, αC, αD, αE, αF, αG, αH, αI) (Figure 1). Helices αA, αB, αD form the dimerization interface, while αC, αE, αG, αI form an amphipathic ligand binding groove^3,5–9^. 14-3-3 proteins recognize a binding motif that contains phosphorylated amino acid residues. Although most binding partners of 14-3-3 have binding motifs containing phosphoserine (pSer)^2,10^, the phosphorylated amino acid residue can be also threonine (pThr). FOXO, forkhead transcription factors^11^, aralkylamine N-acetyltransferase (AANAT)^12^, and CDC25A^13,14^ are some examples of 14-3-3 binding partners with pThr in their binding motifs. Yaffe and colleagues^2^ used a pSer-oriented library to identify two consensus binding motifs, RSXpSXP and RXY/FxpSXP that bind to all 14-3-3 isoforms. X denotes any type of amino acid residue, and pS is pSer.

**Figure 1.**
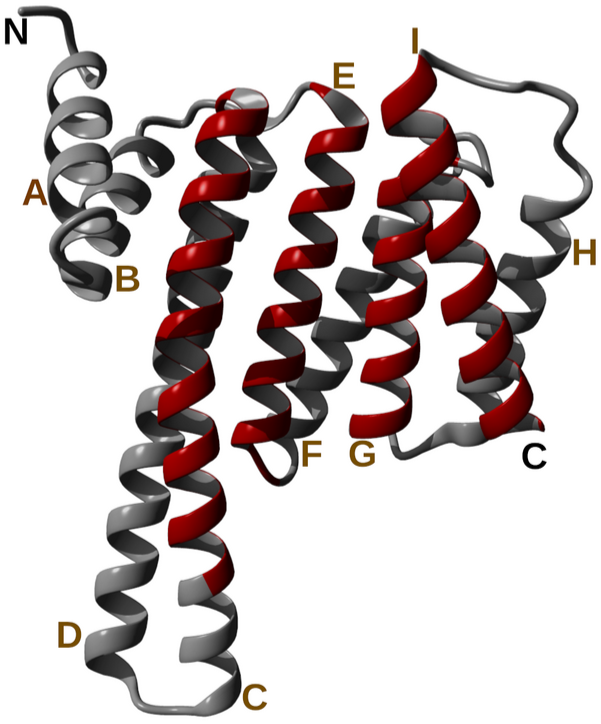
Ribbon representation of monomeric structure of 14-3-3ε. Helices are labeled (A-I) from N- to C- terminal, and helices that form the binding groove are highlighted in red.

The 14-3-3 proteins bind a plethora of proteins, thereby modulating cellular activities, such as, cell cycle control, and apoptosis^11,13–15^. 14-3-3 proteins are associated with various pathological conditions, including neurodegerative disorders like Parkinson’s disease and Alzheimer’s disease^16^. Furthermore, the 14-3-3 proteins are linked to various cancers, such as, glioblastoma ^17^, lung^18^, head and neck^19^, breast^20^ and gastric^21^ cancers. 14-3-3 proteins regulate oncoproteins and tumor suppressor proteins^17–21^. Studies have shown that the 14-3-3ε isoform is involved in various types of malignancies by increasing cell proliferation, invasion or inhibits apoptosis^1,22^. Likewise, increased expression of 14-3-3ε, and its mislocalization from nucleus to cytoplasm, has been reported in cSCC, where it is associated with resistance to apoptosis^14^. In cSCC, 14-3-3ε forms heterodimers with γ and ζ isoforms^23^. Similar to 14-3-3ε, the cell division cycle 25 A (CDC25A) is overexpressed in cSCC and inhibits apoptosis. This anti apoptotic activity of CDC25A is dependent on its binding to a 14-3-3ε heterodimer at the region of pSer178 and/or pThr507 residues^14,23,24^. Thus, inhibition of the 14-3-3ε - CDC25A interaction is a promising target for development of therapeutics for cSCC.

Efforts have been taken previously to control 14-3-3 – ligand binding using different strategies, which includes use of 14-3-3 - binding defective mutant, which negatively affects 14-3-3 mediated processes^25^. Moreover, peptide inhibitors of 14-3-3, such as, R18 and difopein have been used in functional studies^26^ to prevent 14-3-3 – ligand binding interactions. R18 is a 14-3-3 ligand isolated from a phage display library screen for peptide ligand for 14-3-3τ, but R18 also binds other 14-3-3 isoforms with high affinity^26^. The 14-3-3 – R18 interaction is not dependent on phosphorylation, although, X-ray structures ^27^ revealed that Trp-Leu-Asp-Leu-Glu (residues 10-14) pentapepide segment of R18 occupies the same amphipathic binding groove of 14-3-3 as a phosphopeptide does, and negatively charged residues Asp^12^ and Glu^14^ establish contacts similar to that of the pSer of phosphopeptides. R18 inhibits the interactions of 14-3-3 and its clients such as exoenzyme S^28^, Raf-1^26^ and apoptosis signal-regulating kinase 1 (ASK1)^25^. Difopein, a dimer of R18 peptide, also inhibits the 14-3-3 – Raf-1 interaction, thus inducing apoptosis^29^. FOBISIN 101, a pSer/Thr mimetic small molecule can block interactions of 14-3-3 with its clients, Akt and Raf-1. FOBISIN 101 also blocks the ability of 14-3-3 to activate exoenzyme S ADP-ribosyltransferase^30^.

MD simulations have been used to study 14-3-3 proteins and characterize their molecular interactions with their binding partners. Nagy and colleagues^7^ revealed conformational changes of 14-3-3ζ in bound and apo states, as well as the binding mechanism of phosphopeptides to 14-3-3ζ. Using enhanced conformational sampling, Higo and colleagues^31^ elucidated the binding process of phosphorylated myeloid leukemia factor 1 peptide to 14-3-3ζ, and identified various interactions required for 14-3-3ζ – phosphopeptide binding. Furthermore, MD simulations of 14-3-3η – phosphopeptide complexes show the effect of phosphorylation on 14-3-3 - ligand interactions, and indicated interactions between the phosphate group of the peptides and basic amino acid residues of 14-3-3η^32^. Andrei and colleagues^33^ also used MD simulation to rationally design high affinity stabilizers of 14-3-3 protein-protein interactions.

In the previous work by this group^24^, a novel,14 amino acid residues peptide was designed which corresponds to one of the two binding regions of CDC25A (Ac-CDC25A(502-515)-NH_2_ (pT)) to 14-3-3ε. The peptide binds to 14-3-3ε, interrupts its interaction with CDC25A, and induces cell death of cSCC cells with an IC_50_ of 22.1 μM. In the current work, we optimize the structure of the pT to improve its affinity for 14-3-3ε. Initially, we truncated the pT peptide to obtain its 9 amino acid residue analog, Ac-CDC25A(502-510)-NH_2_ (pT(502–510)). Then we modified the pT(502-510) in position 510 to improve its binding affinity for 14-3-3ε. Here, we report peptide analogs that have higher binding affinities for 14-3-3ε than the original pT peptide.

## 2. MATERIALS AND METHODS

### 2.1. Molecular Dynamic Simulations

#### 2.1.1. Structure Preparation

The 14-3-3ε – peptide complexes were generated from the previously studied 14-3-3ε - pT complex^24^. The 14 amino acid residues pT peptide was sequentially truncated to a 9 amino acid residue peptide pT(502-510) using the YASARA program^34^ (Table S1). All the designed peptides were *N*-terminally acetyl protected and *C-*terminally amide protected to maintain the electronic structure of the backbone as it is in CDC25A.

#### 2.1.2. MD Simulations of 14-3-3ε – peptide Complexes

MD simulations were carried out as described in^24^. GROMACS-2021 software package^35^ was used with CHARMM36m forcefield^36^. The 14-3-3ε - peptide complex was solvated in TIP3 water molecules^37^ in a rhombic dodecahedron with the minimal distance of the complex from the edge of the dodecahedron being 1.2 nm. The system was neutralized with Na^+^ and Cl^-^ ions and then the final concentration of NaCl was set to 150 mM. Then the system was subjected to 1000 steps steepest descent energy minimization followed by a 1 ns all atoms positionally restrained constant number of molecules, volume and temperature (NVT) simulation at 300 K using the velocity rescale method^38^, the temperature was kept constant using MD integrator with time constant (τ_T_) of 0.1 ps, and integration time of 2 fs. Next, to ensure the system equilibration and relaxation, the system was subjected to 10 ns unrestrained simulation at a constant number of molecules, pressure and temperature (NPT) at 300 K temperature and 1 bar pressure using the Berendsen methods^39^. With 2 fs integration step, the Lincs algorithm^40^ was used to constrain the hydrogen atoms to their correct length. The system was coupled to constant pressure by the Berendsen method^39^ with isothermal compressibility of 4.5*10^-5^ bar, and to a 300 K temperature by velocity-rescaling method with stochastic term^38^. The time constant for pressure coupling (τ_p_) and temperature coupling (τ_p_) were 0.1 ps and 2 ps, respectively. To calculate the van der Waals interactions, the short-range cutoff was 1.0 nm and long-range cutoff was 1.2 nm using force switch modifier. The long-range electrostatic interactions were calculated using the PME method with 1.2 nm cutoff distance with Verlet scheme and 0.12 nm Fourier spacing. After equilibration, a 500 ns NPT simulation was performed with similar parameters as during equilibration except that the system was coupled to a pressure using the Parrinello-Rahman barostat method^41^ and the temperature was kept constant using stochastic dynamic integrator with, τ_p_ and τ_p_ of 2 ps and 4 ps, respectively, and integration time of 2 fs

#### 2.1.3. Trajectory Analysis

To determine whether the system reached equilibration, configurational entropy was calculated^42,43^. The *covar* module of GROMACS was used to calculate covariance matrix of the Cα-atoms movement, eigenvectors corresponding to the 150 highest eigenvalues were used to calculate configurational entropy as function of time^43^. To determine structural stability, the *rms* module of GROMACS was used to calculate C_α_ atoms root mean square deviation (RMSD) for protein and the docked peptide^44^. The RMSD was calculated for the entire simulation trajectory with reference to the initial structure. Changes in secondary structures of 14-3-3ε and peptides during simulations were analyzed using the defined secondary structure of proteins (DSSP) method^45^. To determine whether aromatic amino acid residues at position 510 of the peptides interact with residues of 14-3-3ε, representative structures of the last 100 ns of the simulations were determined using the GROMOS method of cluster analysis^44^ with C_α_ atom RMSD cutoff of 0.12 nm. Then, using the representative structure, *distance* module of GROMACS was used to identify residues of 14-3-3ε that can interact with aromatic residues in position 510 of the peptides. Distance was calculated between center of mass (COM) of side chain of residues of protein and COM of the ring of aromatic amino acid residue in position 510 of the peptides. 0.8 nm was used as the cutoff distance for interactions^46,47^

#### 2.1.4. Modifying pT(502-510) Using FoldX

To generate various analogs of pT(502-510) peptide, FoldX^48,49^ extension of the YASARA software was used to introduce modifications in the peptide. Here, five representative structures of 14-3-3ε - pT(502-510) complexes were obtained from the most populated clusters of five independent MD simulations. Guided by binding motifs of 14-3-3 proteins, modifications were made in the sequence of the pT(502-510) in each structure by substituting amino acid residue of the peptide with different amino acid residues, one modification at a time (Table S2). The modifications were carried out at 310 K and pH 7.0. Change in free energy of binding (ΔΔG_b_) of 14-3-3ε to peptide due to modifications in the peptide was calculate using FoldX^49^.

#### 2.1.5. Steered Molecular Dynamics (SMD) Simulations of 14-3-3ε – peptide Complexes

For each amino acid modification, the structure that showed the most negative ΔΔG_b_, obtained by FoldX analysis, was used for SMD simulations. The scheme of the SMD simulations is shown in Figure S4. 14-3-3ε - peptide complexes were solvated in TIP3 water molecules in a triclinic box with x, y, and z dimension of 8.5 nm, 9 nm and 13 nm, respectively. All the steps were carried out as described in MD simulations section above, except that, the system was equilibrated in a 100 ps NPT simulation so that the postion of Cα-atoms are restrained in both protein and peptide with 400 kJ mol^-1^ nm^-2^ force constant. Then, the equilibrated system was subjected to a 4 ns steered molecular dynamics (SMD) simulations^50^. Here, position of heavy atoms of 14-3-3ε was fixed, and COM of peptide was pulled in z direction (Figure S4) at a constant velocity of 1.0 nm ns^-1^ and a spring constant of 2500 kJ mol^-1^ nm ^-1^. COM of peptide was defined as the COM of the phosphorylated amino acid residue (pThr^507^), while COM of 14-3-3ε was defined as COM of Lys^50^, Arg^57^, Arg^130^ and Tyr^131^ residues^24^ that interact with pThr^507^.

#### 2.1.6 Umbrella Sampling (US) Simulation

US simulations ^50,51^ were used to determine free energy of binding (ΔG_b_) of the peptides to 14-3-3ε. Snapshots (24-27) from SMD simulations were used as starting configurations for US sampling. The snapshots were taken at every 0.1 nm of the COM distance between the protein and peptide (ξ reaction coordinate). A 4500 kJ mol^-1^ nm^-2^ umbrella potential was applied and each window was simulated for 30 ns. Other parameters used here are the same as in SMD simulations. Trajectories from the US simulation of each window were subjected to the weighted histogram analysis method (WHAM)^52,53^ to obtain potential of mean force (PMF). ΔG_b_ was determined from the PMF as the difference between the largest and the smallest PMF values. Each PMF curve (Figure 4D) was obtained from a combination of histograms from five independent simulations during WHAM analyses. The errors were evaluated over 100 rounds of bootstrap analysis.

### 2.2. Protein

(His)_6_-14-3-3ε was commercially obtained from Novus Biologicals (Centennial CO, USA), and 14-3-3ε-(His)_12_ was expressed in our laboratory (see details on protein expression in the supporting information). The 14-3-3ε was expressed as a *C*-terminal His_12_-tagged protein in E. *coli* BL21 (DE3) by induction with IPTG. The cells were harvested and lysed by sonication in lysis buffer (20 mM Tris, 200 mM NaCl, 20 mM imidazole, 1mM DTT, pH 7.4), the cell lysate was centrifuged and the supernatant was concentrated using a 10,000 Da MWCO Amicon spin filter by centrifuging at 4,000 rpm at 4 °C. The protein was initially purified by immobilized metal affinity chromatography (IMAC) using a HisTrap column (GE HelathCare, HisTrap FF Crude histidine-tagged protein purification columns). The column was equilibrated with 20 mM Tris, 200 mM NaCl, 20 mM imidazole, 1mM DTT, pH 7.4 buffer. The protein was eluted with an imidazole gradient of 20 mM to 500 mM at a flow rate of 0.5 mL/min. The pooled fractions were concentrated into 20 mM Tris, 150 NaCl, 1 mM DTT, pH of 7.4 buffer using 10,000 Da MWCO Amicon spin filter and centrifuging at 4,000 rpm at 4 °C. The protein was further purified by size exclusion chromatography (SEC) using a superdex 75 column (Cytiva, superdex 75 increase 10/300 GL). The protein was eluted with the same buffer at a flow late of 0.5 mL/min. The purified protein was stored at -80 °C.

### 2.3. Peptides

Peptides were either synthesized or commercially obtained from Biosynth International Inc, (Louisville KY, USA). All N-α-Fmoc-protected amino acids were obtained from CEM Corporation (Matthews NC, USA). Peptides were synthesized using standard Fmoc chemistry with Bzl, Boc, tBu, Pbf side chain protections. The peptides were synthesized in 0.1 mmol scale on Rink amide resin using CEM Discover microwave peptide synthesizer. Fmoc deprotection was performed using 20% piperidine in DMF at for 2 minutes and couplings were performed using DIC/Oxyma pure for 3 mintutes at 75 °C. Single coupling cycles were performed for all amino acids, expect double coupling for pThr. Peptides were cleaved from resin and simultaneously deprotected by stirring it in a cocktail containing TFA/Thioanisol/TIPS/H_2_O/DODT in 85:5:2.5:2.5:5 (v/v/v/v/v) for 1 h on ice, and then 2 h at room temperature. The resin was then removed by filtration and the peptide was precipitated with ice-cold diethyl ether and collected by filtration. The crude peptides were dissolved in 10% AcOH in H_2_O, and lyophilized. The crude products were purified by semi-preparative reversed-phase HPLC using C18 column (Phenomenex, Jupiter 300 Å, 5 μm, 250 × 10 mm). The peptides were eluted using 0.1% TFA aqueous solution (v/v, solvent A), and 0.09% TFA in acetonitrile (v/v, solvent B) with a linear gradient of solvent B of 3 to 60 % over 60 minutes, at a flow rate of 4 mL/min. Purity of the peptides were analyzed by analytical HPLC using C18 (Jupiter 300Å 5 μm, 250 × 4.6 mm) column, with the same gradient of solvent B over 40 minutes, at a flow rate of 1mL/min. Masses of the peptides were determined by MALD-TOF mass spectrometry (Bruker, Billerica MA, USA).

### 2.4. Electronic Circular Dichroism (ECD) Spectropolarimetry

Peptides were dissolved in either nanopure water or 30%, 50%, and 75 % (v/v) 2,2,2-trifluoroethanol (TFE) in water. Using Jasco J-810 spectropolarimeter (Jasco Inc., Easton, MD), ECD spectra of the peptides were measured at 20 °C, in a quartz cell with a 0.05 cm pathlength. ECD spectra of each peptide were obtained from average of 20 scans from 180 nm to 260 nm at a rate of 100 nm/min. Mean residue ellipticity was calculated using peptide concentration determine using NanoDrop One (Thermo Scientific) spectrophotometer at 280 nm wavelength. To quantify secondary structure of the peptides, the ECD spectra were deconvoluted using. DichroWeb CD analysis web server and the CDSSTR method with reference set 6 ^54,55^.

### 2.5. Differential Scanning Fluorimetry (DSF)

Melting temperature (T_m_) of 14-3-3ε - peptide complexes were determined by DSF using the BioRad CFX384 Touch real time PCR (Hercules CA, USA). The protein unfolding was detected with fluorescent SYPRO-orange dye (Invitrogen). The assay was performed in 20 mM Tris, 150 mM NaCl, 1 mM DTT PH7.4 buffer, in a 386 well plate with a total volume of 20 μl, containing 2 μM and 30 μM, final concentrations of protein and peptide, respectively, and 500x dilution of the SYPRO-Orange dye stock from the manufacturer. Excitation filter for the dye was 450-490 nm and the emission filter was 560-580 nm. The temperature was held at 25 °C for 5 min, then increased from 25 °C to 95 °C at a rate of 0.3 °C/min, and fluorescence was recorded over time. T_m_ values were determined by plotting the first derivative of the melting curve using GraphPad Prism software.

### 2.6. Surface Plasmon Resonance (SPR)

Binding affinities of pT peptide analogs for 14-3-3ε were measured by SPR using Biacore 8K Surface plasmon resonance instrument (Cytiva, Marlborough MA, USA). 100 nM of 14-3-3ε-(His)_12_ protein in 20 mM Tris, 150 mM NaCl, 50 μM EDTA, 0.005% Tween 20, pH 7.4 buffer was immobilized to a NiCl_2_ activated NTA chip (Cytiva, Marlborough MA, USA) at a flow rate of 5 μL/min for 10 min, and then washed with the running buffer for 20 min. Then peptide solutions at concentrations range of 1 nM-2000 nM were injected cycle-wise at the flow rate of 30 μL/min for 1 min, and dissociation time was set for 3 min. All SPR experiments were performed at 25 °C. The affinities and kinetics of the peptides binding to 14-3-3ε were determined by steady state and 1:1 binding models, respectively, using Biacore Insight Evaluation Software (version 5.0.18.22102). At least 3 replicates were done for each peptide at similar condition, binding affinities (K_D_) are presented as mean ± SD. K_D_ values were used to calculate binding free energy (ΔG) with the following formula: ΔG = RT ln K_D_ (R =1.986 cal mol^-1^ K^-1^, T = 298K).

## 3. RESULTS

### 3.1. Rational Design of pT Peptide Analogs

The Ac-CDC25A(502-515)-NH_2_ (pT) binds to 14-3-3ε and induces death of cSCC cells with an IC_50_ of 22.1 μM^24^. Thus, here we optimized the peptide structure to improve its binding affinity to 14-3-3ε. The proposed binding motifs of 14-3-3 proteins are defined by six to seven amino acid residues, RSXpSXP and RXY/FXpSXP^2^. Furthermore, X-ray structures of 14-3-3 in complex with various ligands showed that 14-3-3 proteins form complexes with short peptides^56^. Hence, to optimize the pT peptide, we sequentially truncated it from both the *N-* and *C-* termini (Table S1). The structures of truncated peptides in complexes with 14-3-3ε were studied using 500 ns MD simulations. During simulations the complexes, protein and peptides in the complexes went through various structural transitions as indicated by the RMSD of Cα atoms (Figure S1), but no peptide dissociation was observed for any peptide analogs. Since the short, 9 amino acid residue analog, pT(502-510) bound to 14-3-3ε (Figure 2A, S2), we used it for further analog design (detailed analysis of the simulation is given below). Using FoldX extension of YASARA program, amino acid residues at different positions of pT(502-510) in the representative structures from five MD simulations (see methods) were replaced with various amino acids (Tables 1 and S2). Binding energies of the resultant peptides with 14-3-3ε were determined. Based on the binding motif of 14-3-3 proteins^2^, and the 14-3-3ε specific binding motif, RF/R/ARpSAPF^10^, either small hydrophobic, positively charged or aromatic residues were introduced at positions that have been shown to be crucial for binding. Substitutions of Gly^510^ with aromatic amino acid residues were the most favorable among the modifications (Table 1). To find out if this is due to aromaticity or hydrophobicity of the residues, we introduced hydrophobic aliphatic amino acid residues Ala and Val in position 510 as well. Although Gly^510^ to Phe modification resulted into the most favorable modification, constraining its side chain x^1^ torsional angle can further improve binding affinity of the peptide. Thus, Phe^510^ was replaced with 1,2,3,4-tetrahydroisoquinoline-3-carboxylic acid (Tic), an analog of Phe that is conformationally constrained in the x^1^ torsional angle rotation ^57^ (Table 1 and Figure S6B). The binding of the designed peptides to 14-3-3ε was further studies using biophysical methods.

**Figure 2.**
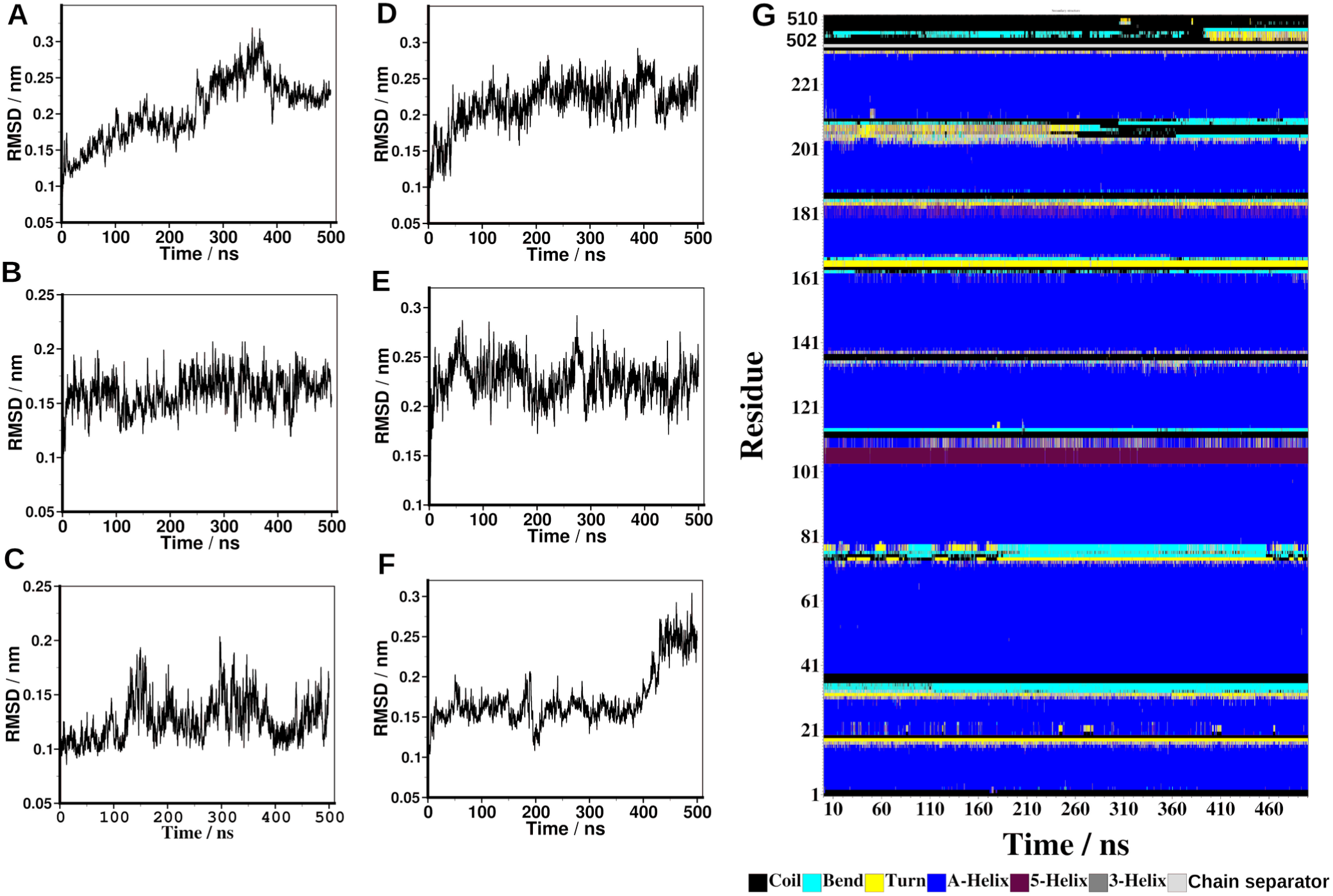
Root mean square deviations (RMSD) of Cα-atoms, and secondary structure of 14-33ε –peptide complexes. 14-3-3ε in complex with: (A) pT(502-510). (B) [Phe^510^]pT(502-510). (C) [Trp^510^]pT(502-510). (D) [Tyr^510^]pT(502-510). (E) [Ala^510^]pT(502-510). (F) [Val^510^]pT(502-510). (G) change in secondary structure of 14-3-3ε – pT(502-510) complex during a 500 ns MD simulation. Secondary structure content was determined the by DSSP method.

**Table 1.** Peptide Analogs of Ac-CDC25A(502-515)-NH_2_ (pT), and the change in binding energies upon substituting residue 510.

| Peptide | Amino acid sequence | $\Delta\Delta G \pm SE / \text{kJ mol}^{-1}$ |
| --- | --- | --- |
| pT | Ac-RTKSRpTWAGEKSKR-NH<sub>2</sub> |  |
| pT(502-510) | Ac-RTKSRpTWAG-NH<sub>2</sub> |  |
| [Phe<sup>510</sup>]pT(502-510) | Ac-RTKSRpTWA<sup>F</sup>-NH<sub>2</sub> | $-5.06 \pm 3.20$ |
| [Trp<sup>510</sup>]pT(502-510) | Ac-RTKSRpTWA<sup>W</sup>-NH<sub>2</sub> | $-2.35 \pm 1.94$ |
| [Tyr<sup>510</sup>]pT(502-510) | Ac-RTKSRpTWA<sup>Y</sup>-NH<sub>2</sub> | $-6.18 \pm 2.77$ |
| [Ala<sup>510</sup>]pT(502-510) | Ac-RTKSRpTWA<sup>A</sup>-NH<sub>2</sub> | $1.45 \pm 0.94$ |
| [Val<sup>510</sup>]pT(502-510) | Ac-RTKSRpTWA<sup>V</sup>-NH<sub>2</sub> | $1.36 \pm 1.61$ |
| [Tic<sup>510</sup>]pT(502-510) | Ac-RTKSRpTWA<sup>Z</sup>-NH<sub>2</sub> | N.D |

Modified residues are highlighted in red. Change in binding energies ( $\Delta\Delta G_b$ ) between 14-3-3 $\epsilon$ – peptide complexes were calculated using FoldX. The error was evaluated over n=5 independent calculations and N.D: not determined.
\*Z, Tic; Ac and -NH<sub>2</sub>, acetyl and amide, respectively, protecting groups.

### 3.2. MD Simulations of the Modified Peptides in Complex With 14-3-3ε

To determine whether the pT(502-510) and its designed analogs form stable complexes with 14-3-3ε, we carried out 500 ns MD simulations. Configurational entropy (Figure S2) shows that, all systems experienced a steep decrease in the first 50 ns, followed by a gradual decrease between 50ns and ∼300 ns, after which the systems reached equilibrium. RMSD of Cα-atoms as a function of time (Figure 2) shows that all peptides formed stable complexes with 14-3-3ε. 14-3-3ε - peptide complexes had RMSD between 0.1 nm and 0.3 nm. Complexes of 14-3-3ε with pT(502-510), [Tyr^510^]pT(502-510), and [Val^510^]pT(502-510) showed gradual increase in RMSD, which became stabilized after 400 ns. Conversely, complexes containing [Phe^510^]pT(502-510), [Trp^510^]pT(502-510), and [Ala^510^]pT(502-510) showed a constant RMSD throughout simulations (Figure 2). RMSDs of 14-3-3ε and the peptide separately, in all complexes, peptides showed larger RMSD than the 14-3-3ε (Figure S2). DSSP analysis of the 14-3-3ε – pT(502-510) complex showed that 14-3-3ε did not experience significant structural changes throughout the simulation. The peptide also maintained its structure with some coil to bent and turn transition of residues 504-506 in the last 150 ns of the simulation (Figure 2G). Both protein and peptides maintained their structures in other protein – peptide complexes as well (Figure S3).

Having determined that aromatic amino acid residues are the most favorable substitutions in position 510 of pT(502-510), we identified amino acid residues of 14-3-3ε that can interact with the aromatic amino acid residues in position 510 of the peptides. The following residues were identified within 0.8 nm of COM of the ring of Phe^510^, Trp^510^ and Tyr^510^ amino acid residue of peptides: Asn^51^, Asn^43^, Ser^46^, Val^47^, Phe^120^, Lys^123^, and Asp^216^ (Figure 3). Phe^510^ had the highest number of interacting residues within 0.8 nm of its COM compared to Trp^510^ and Tyr^510^ (Figure 3). The different types of amino acid residues of 14-3-3ε that were identified interact with the aromatic residues of the peptides through π-π, cation/anion-π, CH-π, and amide-π interactions^46,47^.

**Figure 3.**
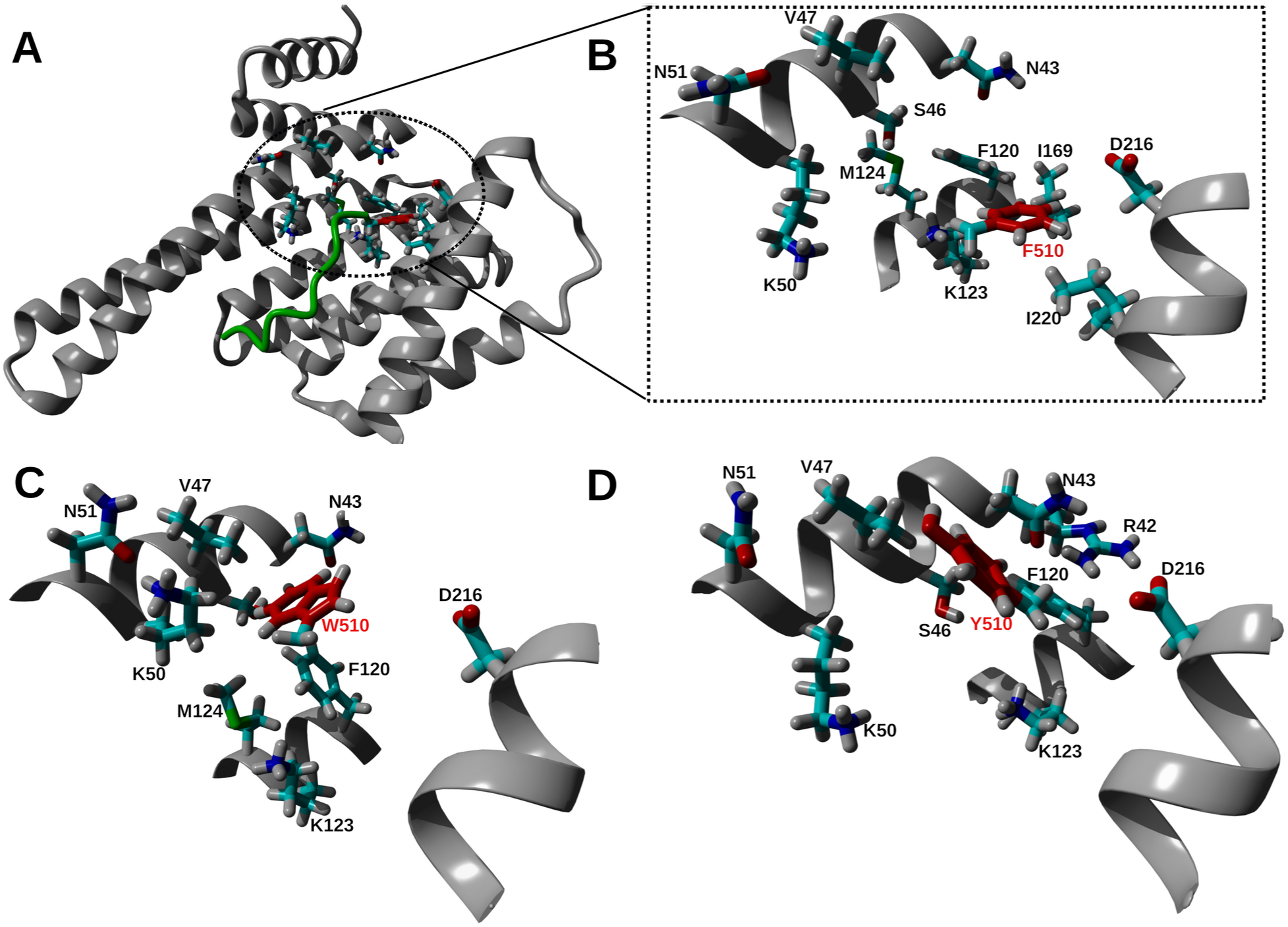
Interacting residues in the 14-3-3ε – peptide complexes. The structures of 14-3-3ε and [Phe^510^] pT(502-510) peptide are shown in gray ribbon and green stick, respectively. Residues of 14-3-3ε that are within 0.8 nm of center of mass (COM) of the ring of aromatic amino acid residues in position 510 of peptides are in colored sticks. Amino acid residues are indicated in one letter code. The aromatic residue in position 510 of the peptide are shown by red sticks. Representative structures from 500 ns simulations were used. (A) The structure of 14-3-3ε – [Phe^510^] pT(502-510) complex showing residues of the protein identified within 0.8 nm from COM of Phe^510^ of the peptide. (B, C and D), a close-up view of the identified amino acid residues when 14-3-3ε is in complex with [Phe^510^] pT(502-510), [Trp^510^] pT(502-510) and. [Tyr^510^]pT(502-510) peptides, respectively.

### 3.3. Binding Free Energy Analysis Using Umbrella Sampling Method

To determine ΔΔG_b_ of modified peptides with 14-3-3ε FoldX used one snapshot from an MD trajectory at a time. Therefore, to further assess binding of the peptides to 14-3-3ε, further MD simulations were performed. We first carried out SDM simulations, then, snapshots from SMD trajectories were subjected to umbrella sampling simulations^50^ and analyzed by the WHAM method^52^. Peptides were pulled away from the binding groove of 14-3-3ε by applying a force with spring constant of 2500 kJ mol^-1^ nm ^-1^ on the COM of pThr507 residue of the peptides at a constant rate of 1 nm/ns. As shown by the force-time curve (Figures 4A and S5), all peptides show rupture event around at 500 ps with rupture force of 700-900 kJ mol^-1^ nm, except, the [Trp^510^]pT(502-510) peptide for which the rupture occurred around at 1000 ps. The distance-time curves show minimal change in distance between COM of 14-3-3ε and COM of peptide in the first 500 ps, and a gradual change over the rest of the simulation time (Figures 4B and S5) indicating a peptide dissociation after rupture. Curves in the US simulation histograms (Figure 4C) are overlapping, indicating that the simulations are well suited for WHAM analysis. The PMF curves (Figure 4D) show that all peptides experienced gradual increase in PMF up to reaction coordinate (ξ) of ∼2 nm, after which the peptide dissociated from the protein, thus, PMF stayed constant. ΔG_b_ derived from PMF is shown in Figure 4E, all peptides favorably bound 14-3-3ε as indicated by negative ΔG_b_.14-3-3ε - pT(502-510) complex has ΔG_b_ of -92.24 ± 6.56 kJ mol^-1^. The 14-3-3ε - [Phe^510^]pT(502-510) formed a complex with the most negative ΔG_b,_ -143.99 ± 6.35 kJ mol^-1^, while 14-3-3ε - [Val^510^]pT(502-510) showed the lowest ΔG_b_ of -80.02 ± 5.47 kJ mol^-1.^ All other analogs formed complexes with ΔG_b_ in the range of -98.16 to -109.80 kJ mol^-1^.

**Figure 4.**
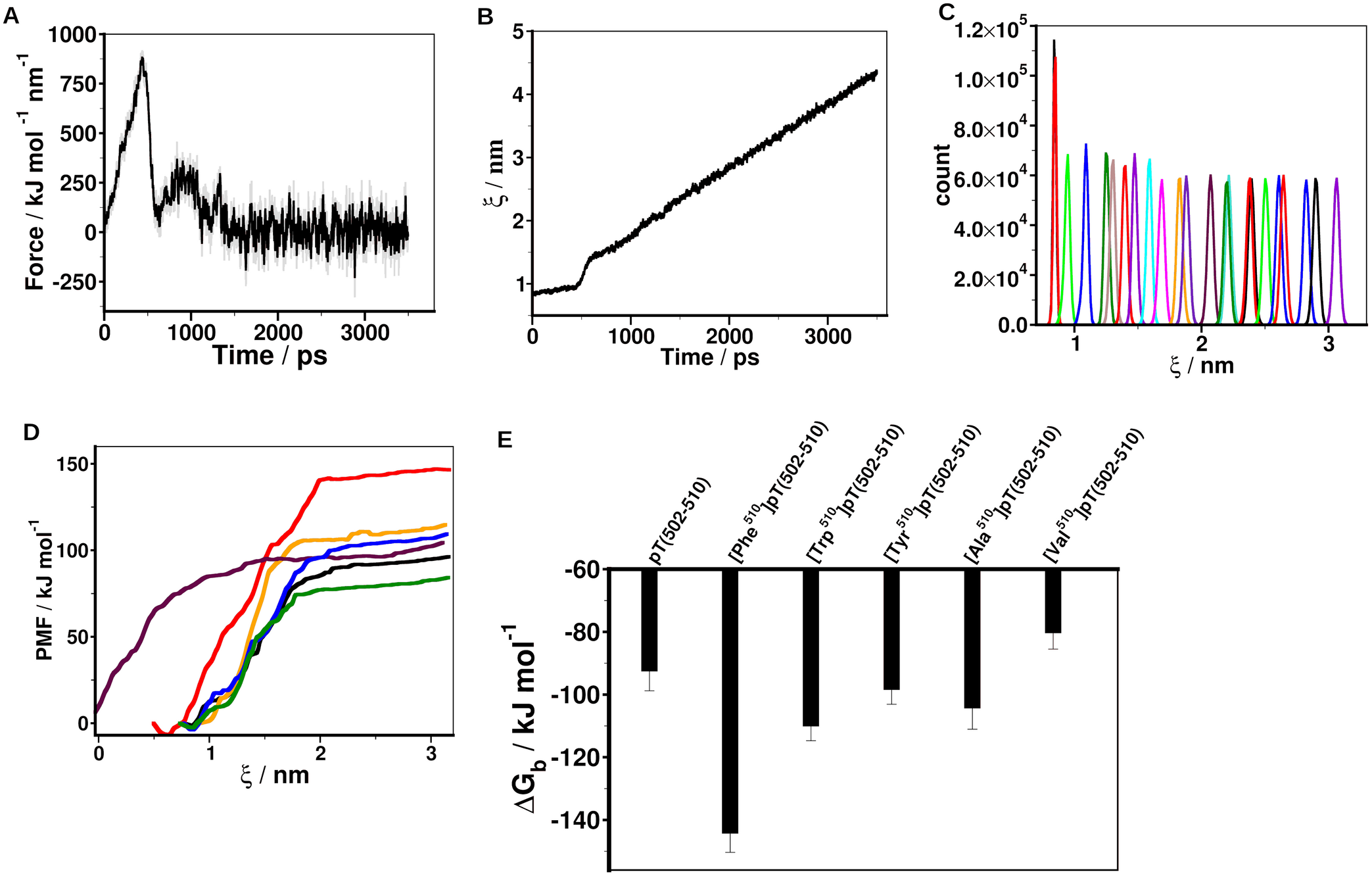
Steered molecular dynamics (SMD) simulations of 14-3-3ε – peptide complexes. (A) Time dependent external force applied to pull pT(502-510) away from the binding groove of 14-3-3ε. (B) Time dependent distance between center of mass (COM) of 14-3-3ε and COM of pT(502-510). (C) Converged umbrella histograms of 27 configurations, each derived from 30 ns simulation of 14-3-3ε - pT(502-510). (D) Potential of mean force (PMF) curves obtained using weighted histogram analysis method (WHAM). pT(502-510) (black), [Phe^510^]pT(502-510) (brown), [Trp^510^]pT(502-510) (orange), [Tyr^510^]pT(502-510) (purple), [Ala^510^]pT(502-510) (cyan), [Val^510^]pT(502-510) (green). (E) Free energy of binding (ΔG_b_) of 14-3-3ε – peptide complexes obtained from potential of mean force (PMF). The free energies of binding were estimated as the difference between the maximum and minimum values of the PMF curves.

### 3.4. Structural Characterization of the Peptides

We used ECD spectropolarimetry to determine secondary structures of the peptide analogs dissolved in water and increasing TFE concentration. In water, the presence of a minimum at ∼198 nm suggests that the peptides adopted an unordered conformation^58^, some of the peptides showed a spectral shift in presence of TFE (Figures 5 and S7). To quantify secondary structure content of the peptides, we analyzed ECD spectra using the CDSSTR method of the DichroWeb CD analysis server. The data (Table S3) shows that secondary structure of the peptides is predominantly random coil with some helicity, β-stand and β-turn conformations. In water, [Tic^510^]pT(502-510) showed the most ordered structure regardless of TFE concentration. The structure of pT(502-510) was not affected by varying concentrations of TFE. Secondary structures of all other peptides became more structured in TFE, especially at 70 % TFE. The parent pT peptide had the highest tendency to assume a more ordered structure with increasing TFE concentration (Figures 5, S7 and Table S3).

**Figure 5.**
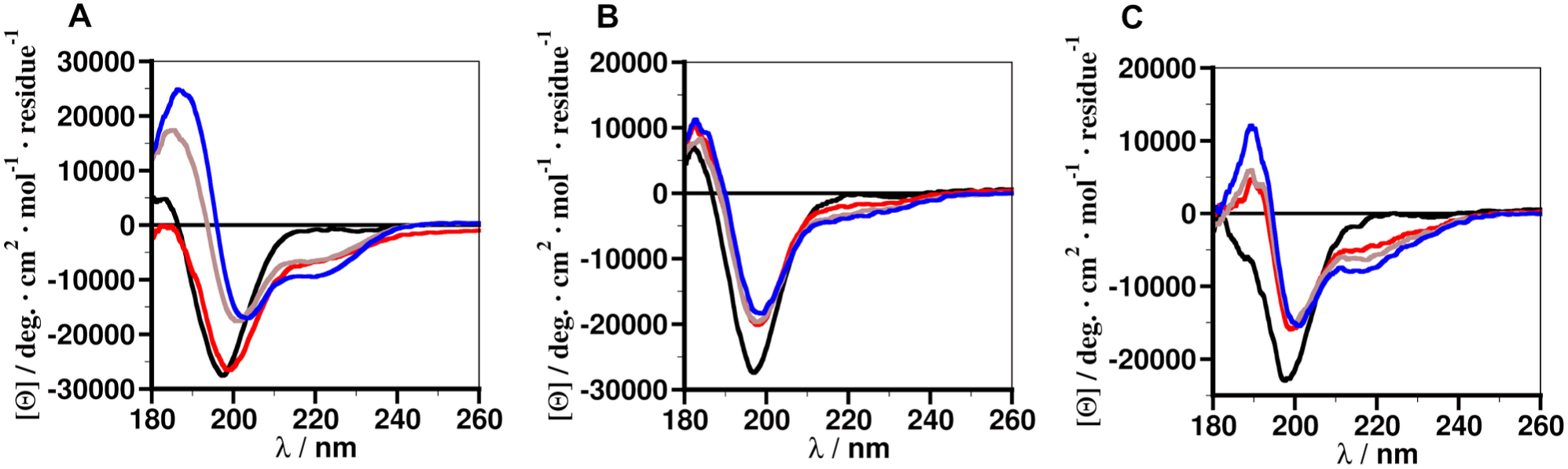
Electronic circular dichroism (ECD) spectra of peptides in different media. (A) pT; (B) pT(502-510); (C) [Phe^510^]pT(502-510). Black, water; red, 30 % TFE; brown, 50 % TFE; blue, 70 % TFE. Each spectrum is an average of 20 scans. Mean residue ellipticity was determined from concentration of the peptides.

### 3.5. Binding of pT Peptide Analogs to 14-3-3ε i*n vitro*

Based on MD simulation results, pT peptide analogs (Table 1) were either synthesized using Fmoc chemistry, or commercially obtained from Biosynth corporation. To confirm our computational results, we deciphered binding of the synthetic peptides to 14-3-3ε by differential scanning fluorimetry (DSF). Figures 6 and S8 show the thermal unfolding profile of 14-3-3ε, and the14-3-3ε - pT(502-510) complex, and their respective derivative curves. DSF results (Table 2) show that all peptides caused a shift in T_m_ (ΔT_m_) of the 14-3-3ε, pT and pT(502-510) peptides induced ΔT_m_ of 2.43 ± 0.12 and 3.70 ± 0.06 °C, respectively. The [Phe^510^]pT(502-510) caused the strongest ΔT_m_ of 5.19 ± 0.41 °C, other analogs led to ΔT_m_ in the range of 3.70-4.44 °C.

**Figure 6.**
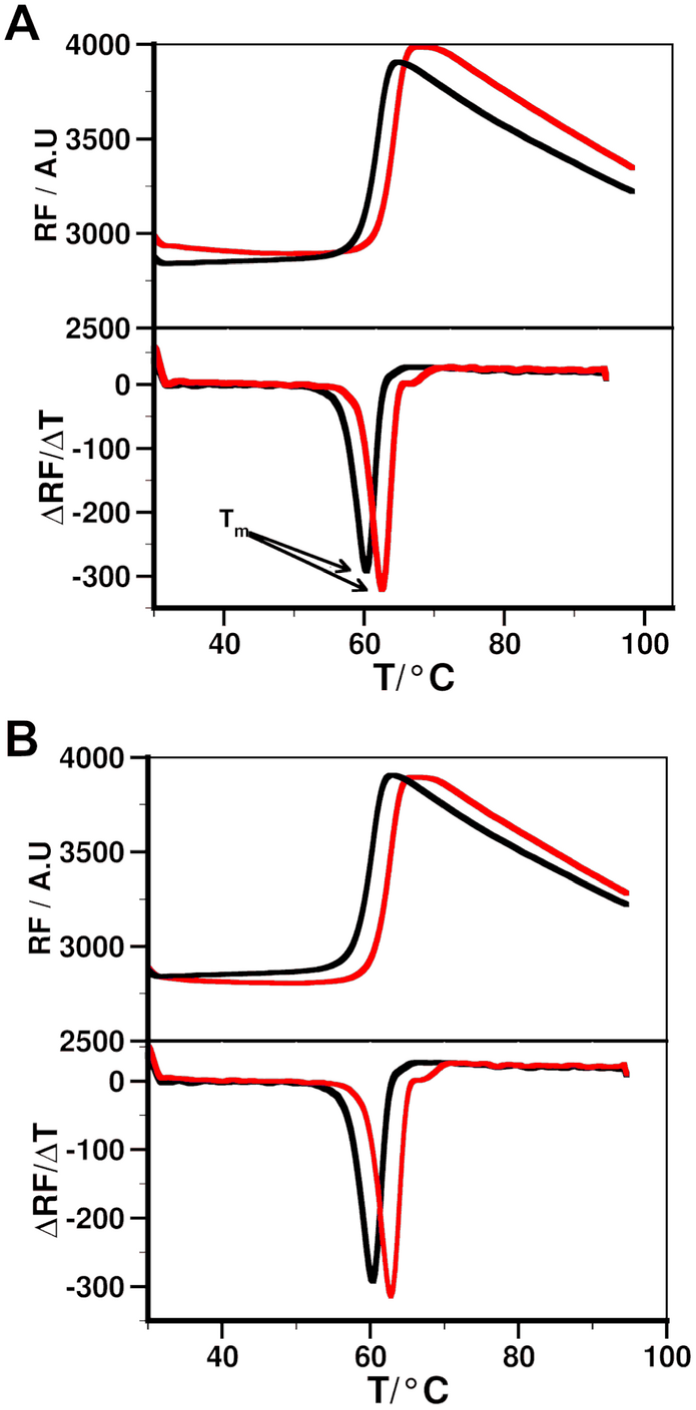
The binding of peptides to 14-3-3ε by differential scanning fluorometry (DSF). (A) Thermal denaturation of 14-3-3ε without (black) and with (red) peptide, and their respective first derivatives, from which melting temperature, T_m_ (inflection point of the curve) is determined; (A) pT; (B) pT(502-510). The presence of the peptide ligand shifts the black curve to the right. The degree of protein stabilization by the peptide can be calculated as change in melting temperature (ΔT_m_) between 14-3-3ε – peptide complex and 14-3-3ε.

**Table 2.** Melting temperature shift and binding affinity of peptide analogs for 14-3-3ε.

| Peptide | $\Delta T_m$ | $K_D \pm SD / \text{nM}$ | $\Delta G \pm SD / \text{kcal mol}^{-1}$ |
| --- | --- | --- | --- |
| pT | $2.43 \pm 0.12$ | $144.33 \pm 2.5$ | $-9.32 \pm 0.01$ |
| pT(502-510) | $3.70 \pm 0.06$ | $45.20 \pm 24$ | $-10.06 \pm 0.29$ |
| [Phe<sup>510</sup>]pT(502-510) | $5.19 \pm 0.41$ | $22.0 \pm 0.77$ | $-10.42 \pm 0.02$ |
| [Trp<sup>510</sup>]pT(502-510) | $3.70 \pm 0.23$ | $76.2 \pm 12.60$ | $-9.71 \pm 0.10$ |
| [Tyr<sup>510</sup>]pT(502-510) | $3.78 \pm 0.32$ | $93.10 \pm 4.57$ | $-9.58 \pm 0.029$ |
| [Ala<sup>510</sup>]pT(502-510) | $4.44 \pm 0.48$ | $75.37 \pm 10.15$ | $-9.71 \pm 0.08$ |
| [Val<sup>510</sup>]pT(502-510) | $3.76 \pm 0.38$ | $211.66 \pm 6.02$ | $-9.12 \pm 0.02$ |
| [Tic<sup>510</sup>]pT(502-510) | $4.43 \pm 0.08$ | $30.85 \pm 0.18$ | $-10.23 \pm 0.02$ |

Change in melting temperature ( $\Delta T_m$ ) due to peptides binding to 14-3-3 $\epsilon$ , and steady state affinities ( $K_D$ ) of peptides for 14-3-3 $\epsilon$ were determined by DSF and SPR, respectively. $\Delta T_m$ values correspond to the average and SD of $n \geq 5$ , and $K_D$ values correspond to the average and SD of $n \geq 3$ . The free energies of binding ( $\Delta G$ ) were calculated from $K_D$ .

To quantify binding affinity of the peptides to 14-3-3ε, we performed SPR and steady state affinities (K_D_) and kinetics of the peptides were determined. Representative sensorgrams are shown in Figures 7 and S8. As shown in Table 2, all the peptides had nanomolar binding affinity for 14-3-3ε, the pT peptide has K_D_ value of 144.33 ± 2.5 nM, while pT(502-510) has K_D_ value of 45.20 ± 24 nM. Kinetics (k_on_ and k_off_) of the peptides are reported in Table S4 (K_D_ values calculated from the kinetics qualitatively reproduce the steady state affinities). Among all pT peptide analogs [Phe^510^]pT(502-510) has the highest affinity for 14-3-3ε with a K_D_ of 22.0 ± 0.77 nM, while [Val^510^]pT(502-510) has the weakest affinity for 14-3-3ε with K_D_ of 211.66 ± 6.02 nM. [Tic^510^]pT(502-510) peptide had the affinity that is comparable to that of [Phe^510^]pT(502-510). Free energy of binding (ΔG) derived from K_D_ shows that all peptides bound 14-3-3ε with negative ΔG. (Table 2).

**Figure 7.**
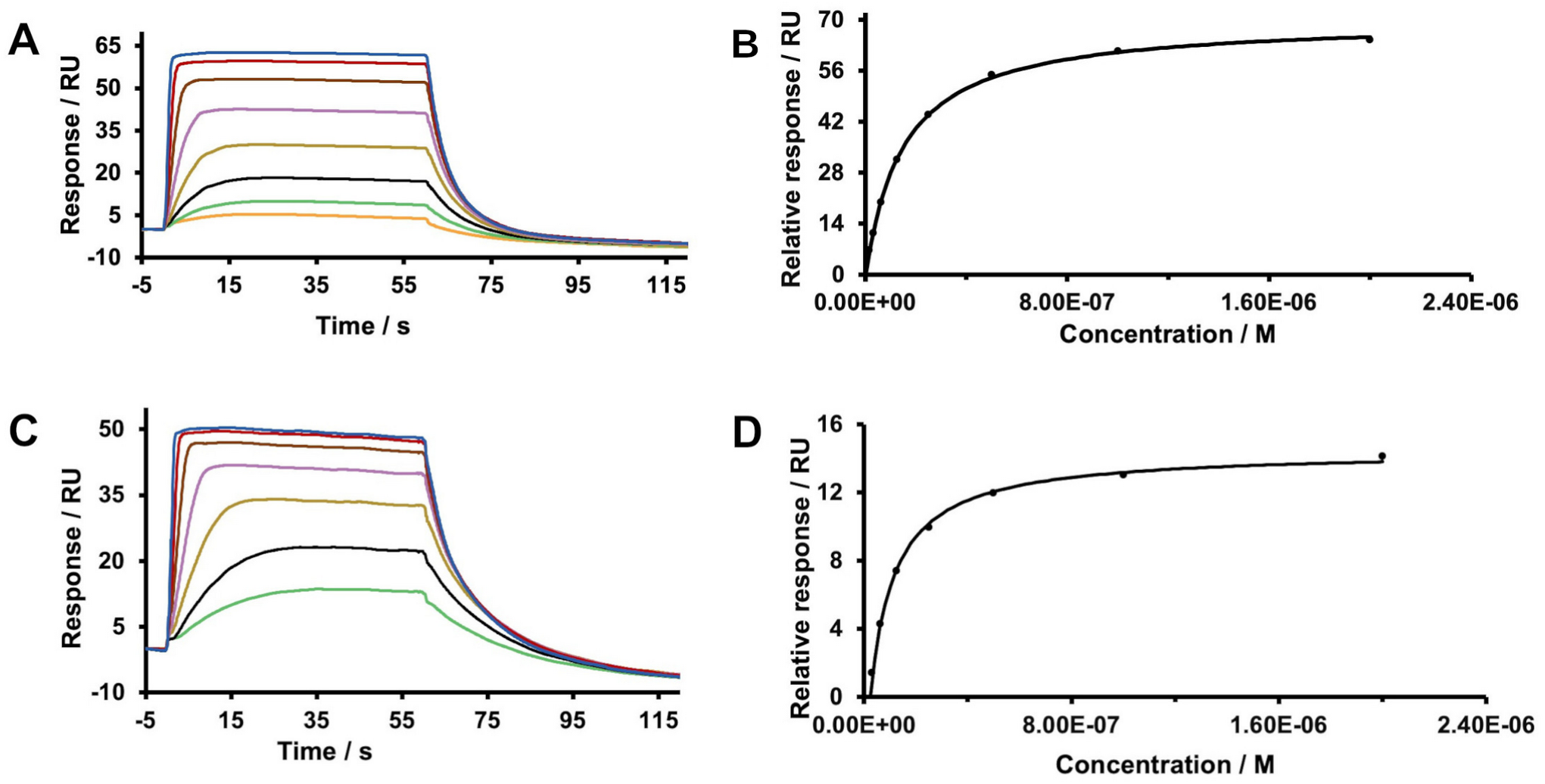
The binding of peptide ligands to 14-3-3ε by surface plasmon resonance (SPR). Sensogram and binding isotherm of 14-3-3ε – pT (A, B) and 14-3-3ε – pT(502-510) (C, D) complexes. 14-3-3ε was immobilized on sensor chip and the binding of peptides at different concentrations (1-2000 nM) were measured at 298 K. Steady state affinities of the peptides (K_D_) were determined using Biacore analysis software.

## 4. DISCUSSION

In this work, by employing an array of different computational and experimental methods, we have shown that shortening and modifying the original pT peptide resulted in peptides with improved affinity for 14-3-3ε (Table 2). 14-3-3ε is overexpressed and mislocalized to the cytoplasm in cSCC, and its interactions with CDC25A suppress apoptosis^14^. The previous study by our group^24^ showed that pT peptide binds 14-3-3ε, inhibits its interaction with CDCC25A, and induced death of cSCC cells at a high concentration. Therefore, we aimed to improve binding affinity of the peptide for 14-3-3ε, thus improving its activity against cSCC.

We designed peptide analogs of pT and studied their binding to 14-3-3ε using MD simulations, DSF and SPR. Our approaches to optimization of the pT peptide was motivated by the fact that the binding motifs of the phospopeptides are defined by five to six amino acid residues, and that short peptide ligands form complexes with 14-3-3ε. Upon sequential truncation (Table S1) of the 14 residue peptide pT, its 9 residues peptide analog pT(501-510) still formed a complex with RMSD (Figure 2A) that is comparable to that of pT^24^. We confirmed binding of the peptide to 14-3-3ε using DSF, a method that is used to determine melting temperature of the protein. DSF relies on extrinsic fluorescence like dyes that allows to monitor melting of protein up on heating protein solution at increasing temperature. Ligand binding causes a shift in melting temperature of the protein^59^. DSF and SPR showed that, compared to pT, pT(502-510) caused a larger ΔT_m_ of 14-3-3ε, and its affinity was ∼3 fold higher than that of pT (Table 2). This improvement in binding affinity may be attributed to the short peptide better fitting in the binding groove of 14-3-3ε than the pT peptide. Thus, we introduced various substitutions in the sequence of pT(502-510) to improve its binding to 14-3-3ε (Table 1 and S2).

To target interaction with 14-3-3ε, we specifically introduced aromatic and positively charged amino acid residues to target interactions such as π-π, anion-π, cation-π, and ionic interactions^46,47^ between the peptide and 14-3-3ε. Substitution of Gly in position 510 of the peptide with aromatic amino acid residues resulted into favorable modifications (Table 1). The free energy of binding (ΔG_b_) obtained by SMD and subsequent US simulations further showed that these modifications resulted in peptides that more favorably bound to 14-3-3ε than does pT(502-510) (Figure 4E). This suggests that aromatic amino acid residues in the peptide analogs could induce weakly polar interactions with amino acid residues of 14-3-3ε (Figure 3). We also introduced Ala and Val at position 510 to ascertain whether the improvement in binding of the peptide analogs containing the aromatic amino acid residues is due to hydrophobicity or aromaticity. The calculated ΔΔG_b_ by FoldX for these substitutions were not favorable (Table 1), which were most likely due to a limited number of structural sampling. Therefore, we performed SMD and subsequent US simulations for the analogs. Results showed that they both favorably bound 14-3-3 ε, but ΔG_b_ of Ala-substituted analog was more negative than that of Val. Furthermore, ΔG_b_ of Ala^510^ containing peptide was comparable to that of the peptides containing aromatic amino acid residues (Figure 4E). These results indicate that aromatic and small hydrophobic amino acid residues are the most suitable in position 510 of the pT(502-510) peptide. Structural features of the peptide analogs determined DSSP analysis of the 500 ns simulations and those obtained by ECD measurements indicate that the peptides upon binding do not undergo major structural transition. This implies that peptide binding to 14-3-3ε is not opposed by conformational entropy change of peptides.

DSF measurements showed that all the designed peptides caused a shift in T_m_ of 14-3-3ε. In all melting curves (Figures 6 and S8) of 14-3-3ε, fluorescence increases with temperature and then decreases at high temperature. The decrease in fluorescence signal at elevated temperature is most likely due to protein aggregation and/or dye dissociation from the protein at high temperatures. As determined by SPR, the [Phe^510^]pT(502-510) peptide has the highest affinity, ∼6.5 fold higher than that of the pT peptide (Table 2). This implies that the aromatic side chain of Phe fits best in position 510 of pT(502-510). Given the flexibility of x^1^ angle of Phe (Figure S6A), the Gly^510^ was substituted with Tic amino acid residue to improve conformational stability of the aromatic side chain of Phe^510^. Currently, no CHARMM36m force field parameters are available for Tic amino acid. Thus, no MD simulations were done for the [Tic^510^]pT(502-510) – 14-3-3ε complex. However, DSF measurement showed that the peptide caused significant increase of T_m_ of 14-3-3ε and the K_D_ (36.66 nM) of the peptide for the protein, determined by SPR, is comparable to that of [Phe^510^]pT(502-510) (22.0 nM). Hence, constraining the χ^1^ torsional angle of the Phe residue did not improved further the binding affinity. Based on these results, all modification introduced in pT(502-510) improved its binding except the Val^510^ modification, and [Phe^510^]pT(502-510) is the best peptide candidate for 14-3-3ε (Table 2). Our results are in agreement with the large-scale screening for peptide binding motifs for 14-3-3 isoforms that showed that Phe is the most favorable amino acid residue in position 510 of peptide motif for 14-3-3ε isoform^10.^

## 5. CONCLUSION

In this study, a combination of *in silico* and biophysical approaches were used to design pT peptide analogs, and study their binding to 14-3-3ε. Since it is known that short peptide ligands bind 14-3-3ε, we shortened the pT peptide from 14 to 9 amino acid residues (pT(502-510)). SMD studies indicated favorable binding of pT(502-510) to 14-3-3ε, and SPR and DSF showed that pT(502-510) is a stronger binder of 14-3-3ε than pT. Aromatic amino acid residues are involved in various types of interaction, thus to further improve binding of pT(502-510), we substituted Gly ^510^ within pT(502-510) with aromatic amino acid residues. SMD and subsequent US simulations indicated that aromatic amino acid substitutions of Gly^510^ residue were the most favorable modifications. The simulation results are agreement with both DSF and SPR measurements. All the studies indicated that the [Phe^510^]pT(502-510) peptide is the best binder of 14-3-3ε. Our study also highlights the importance of aromatic amino acid residues in interactions of our peptides with 14-3-3ε. Overall, we have designed peptides that bind 14-3-3ε with nanomolar affinities. The ability of the optimized peptides to induce apoptosis of cSCC cells will be determined in future studies.

## Acknowledgments

This work was supported by the National Institutes of Health R01 CA253573-01 and the state of Nebraska LB595 grants. Mass spectral analysis of synthetic peptides were performed by the Mass Spectrometry Core facility as a component of the Auditory Vestibular Technology Core within the Translational Hearing Center at Creighton University, School of Medicien funded by CoBRE Award 5P20GM139762 from the NIH.

## Notes

### Competing Interest Statement

The authors have declared no competing interest.

