## Supporting Material for "Optimizing Phosphopeptide Structures That Target 14-3-3ε in Cutaneous Squamous Cell Carcinoma"

### METHODS

#### Protein Expression

The 14-3-3ε-$(His)_{12}$ plasmid was purchased from GenScript (Piscataway NJ, USA), and was transformed into BL21(DE3) cells (Invitrogen) according to the protocol provided by the manufacturer. Colonies were grown on LB agar plates containing 50 μg/mL kanamycin for overnight at 37 °C. On the following day, one colony was selected and cultured overnight at 37 °C in 25 mL LB medium containing 50 μg/mL kanamycin. Then 5 mL of the precultured cells was inoculated into 1 L LB media containing 50 μg/mL kanamycin and incubated at 37 °C in a shaking incubator at 200 rpm. When the optical density of the culture at 600 nm wavelength reached around 0.6, 0.5 mM isopropyl ß-D-1-thiogalactopyranoside (IPTG) was added to media to induce 14-3-3ε protein expression. After induction, cell culture was incubated for 3 hours at 37 °C in a shaking incubator. Cells were then harvested by centrifugation at 10,000 x g for 15 min at 4°C. After centrifugation, the cells were left on dry ice for 10 minutes, and then stored at -80°C until further use. For purification, cells were thawed on ice, and then re-suspended in lysis buffer (20 mM Tris, 200 mM NaCl, 20 mM imidazole, 1mM DTT, pH 7.4,) containing 50 mg/mL DNase, 1 mM DTT, 1 mg/mL lysozyme, 0.5% tween 20, and 1 tablet of protease inhibitor (ThermoFisher, Pierce protease inhibitor mini tablets). The cells were lysed by sonication, and cell lysate was centrifuged at 20000 g for 20 minutes. The supernatant was concentrated using a 10,000 Da MWCO Amicon spin filter by centrifuging at 4,000 rpm at 4°C. The protein was purified as described in the method section.

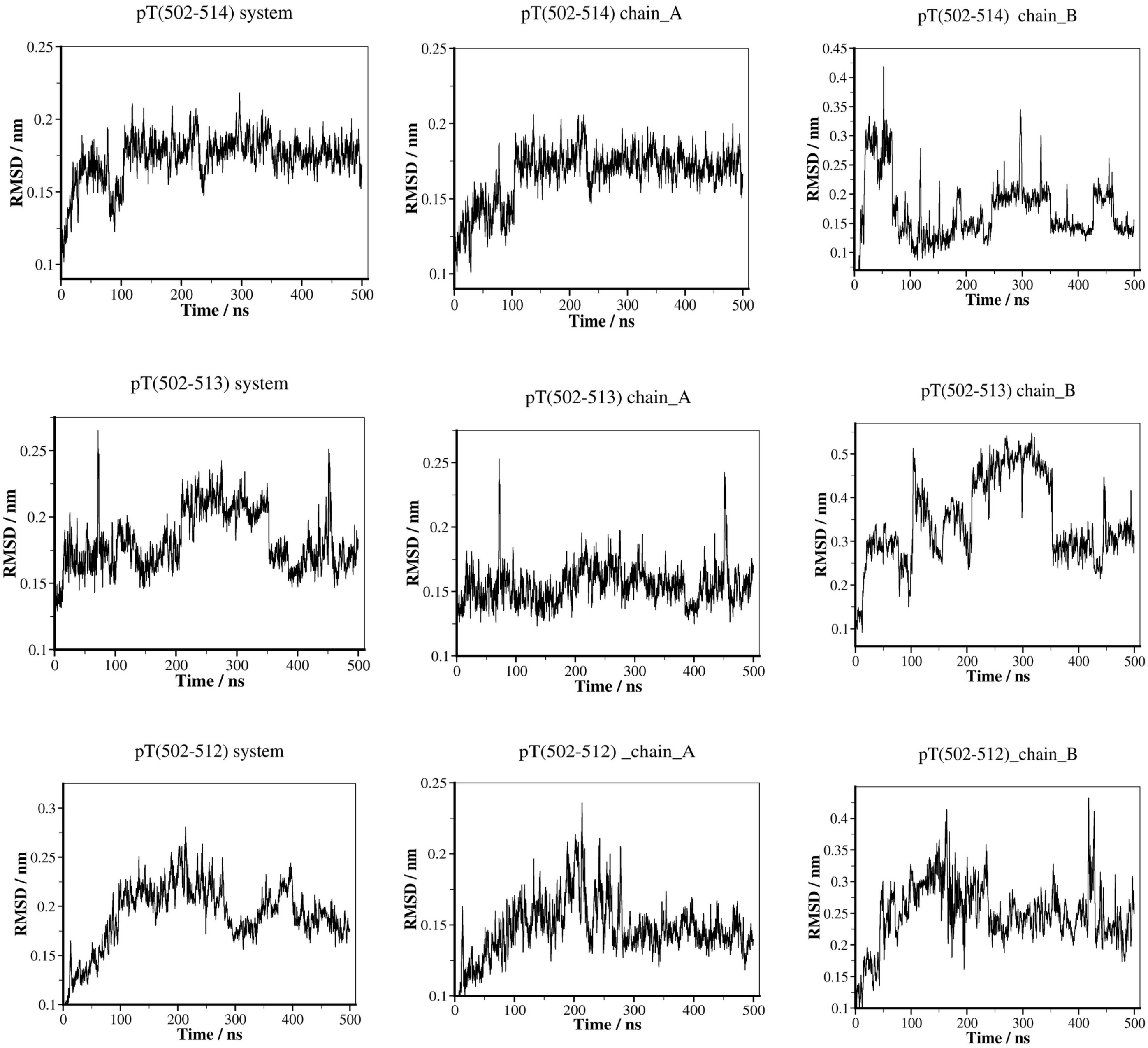
pT(502-514) system
pT(502-514) chain_A
pT(502-514) chain_B
pT(502-513) system
pT(502-513) chain_A
pT(502-513) chain_B
pT(502-512) system
pT(502-512) _chain_A
pT(502-512)_chain_B
RMSD / nm
Time / ns

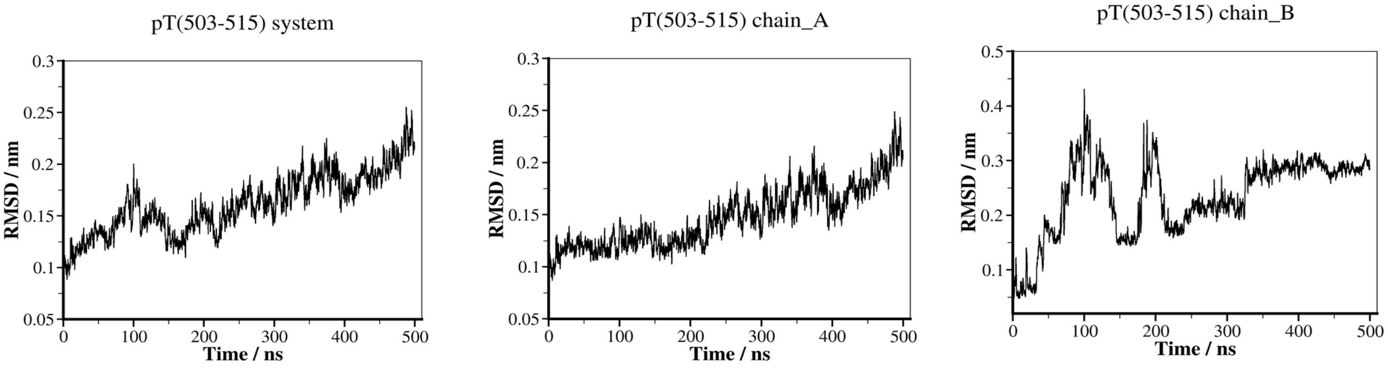


**Figure S1**. **Molecular dynamics simulations of 14-3-3ε – peptide complexes.** A time course of root mean square deviation (RMSD) during simulations of 14-3-3ε in complex with peptide fragments of different lengths (9-13 amino acid residues), system, 14-3-3ε-peptide complex; chain_A, 14-3-3ε; chain_B, peptide. All peptides formed stable complexes with 14-3-3ε, and they did not dissociate from the protein. 14-3-3ε – peptide complex (left panel), 14-3-3ε (middle panel), peptide (right panel).

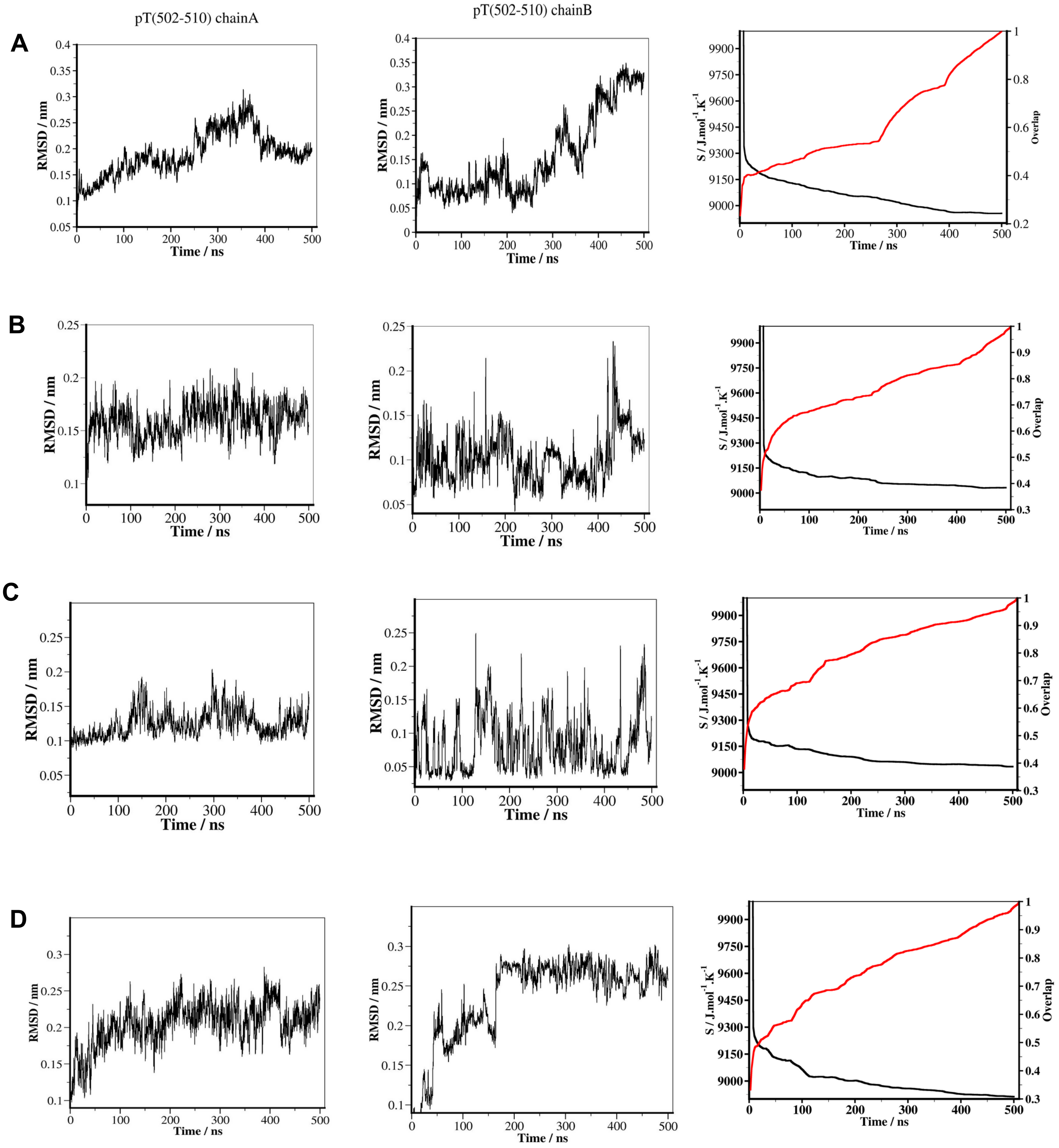
pT(502-510) chainA
pT(502-510) chainB
A
B
C
D
RMSD / nm
Time / ns
S / J.mol$^{-1}$.K$^{-1}$
Overlap

E

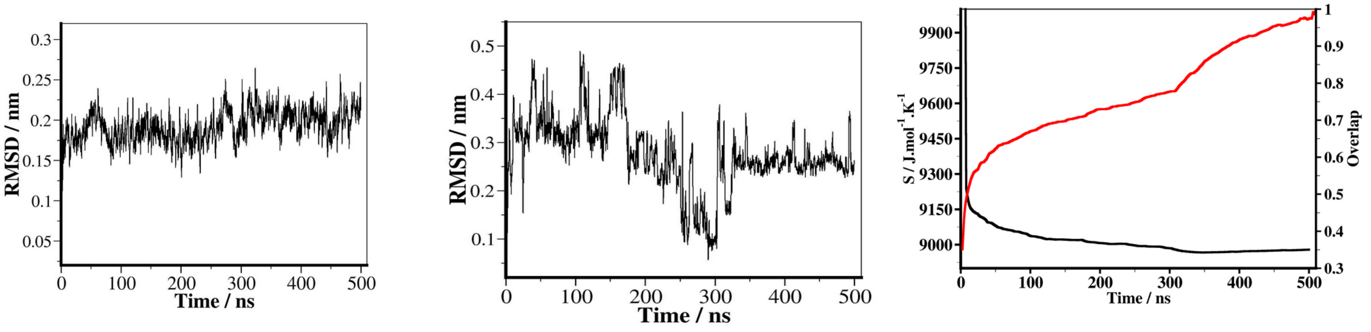


F

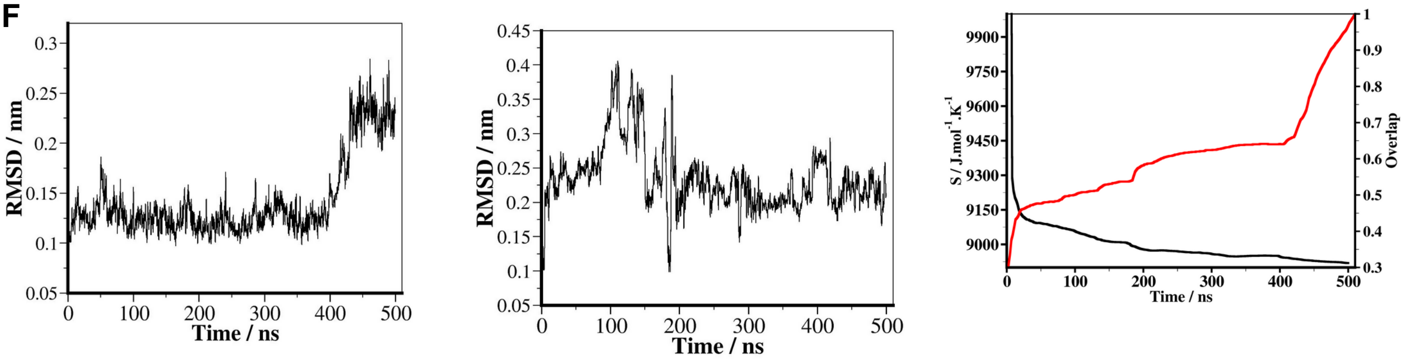


**Figure S2**. Molecular dynamics simulations of 14-3-3ε – peptide complexes. A time course of the RMSD of Cα-atoms, and entropy (black curve) and convergence (red curve) of systems during simulations of 14-3-3ε in complex with the 9 amino acid residues peptide analogs. Protein (left panel); peptide (middle panel); configurational entropy (black) and overlap of sampled region of subspace (red) of system (right panel). (A) pT(502-510). (B) [Phe$^{510}$]pT(502-510). (C) [Trp$^{510}$]pT(502-510). (D) [Tyr$^{510}$]pT(502-510). (E) [Ala$^{510}$]pT(502-510). (F) [Val$^{510}$]pT(502-510). Configurational entropy shows that the system experienced a sudden decrease in the fist 50 ns, then a gradual decrease between 50 ns and ~300 ns, after which the system stabilized. The overlap of the sampled region of subspace for the last 100 ns of simulations is generally greater 0.6 which further support the convergence of simulations[1]. All the peptides formed stable complexes with 14-3-3ε.

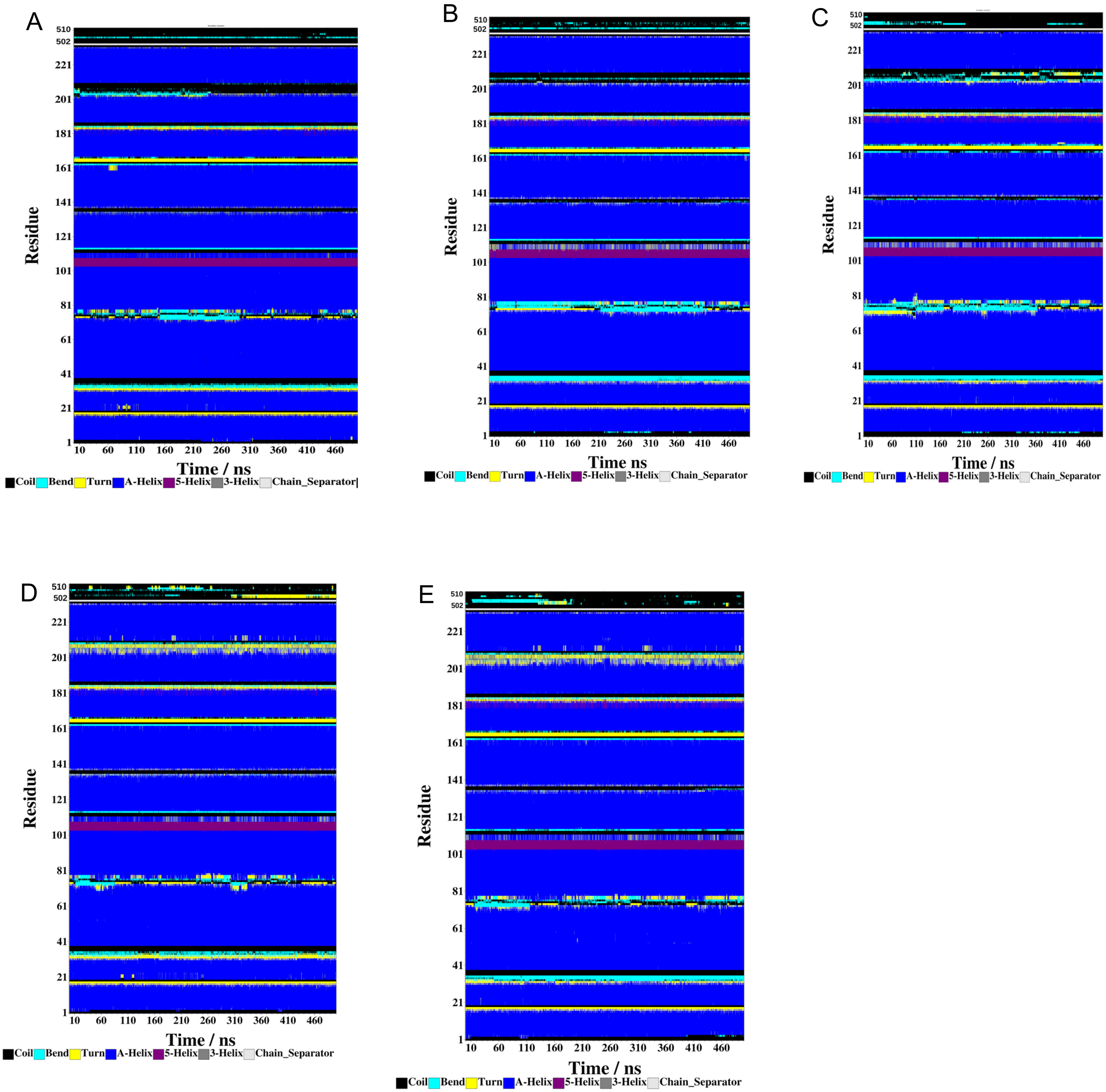


**Figure S3.** Change in secondary structure of 14-3-3ε – peptide complex during MD simulations. (A) [Phe$^{510}$]pT(502-510); (B) [Trp$^{510}$]pT(502-510); (C) [Tyr$^{510}$]pT(502-510); (D) [Ala$^{510}$]pT(502-510); (E) [Val$^{510}$]pT(502-510). Secondary structure content was determined using the defined secondary structure of proteins (DSSP) method. The legend applies to all panels

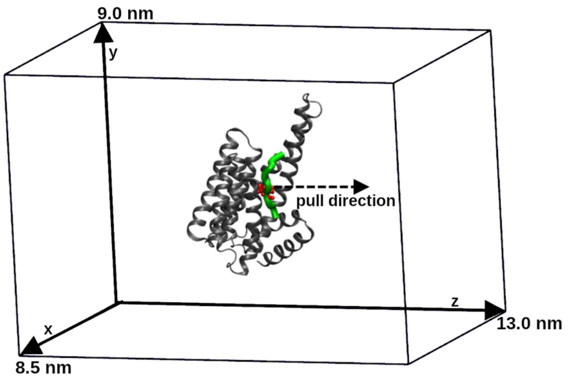


**Figure S4**. Steered molecular dynamics (SMD) simulations set up. 14-3-3ε – peptide complex is in a simulation box, and pulling force is applied on the center of mass (COM) of pThr507 residue of the peptide along the z-axis. 14-3-3ε is shown in gray, the peptide in green, and the pTh507 residue is in red. For clarity, the water molecules are not shown. The dimension of the box is 8.5 nm × 9.0 nm × 13 nm.

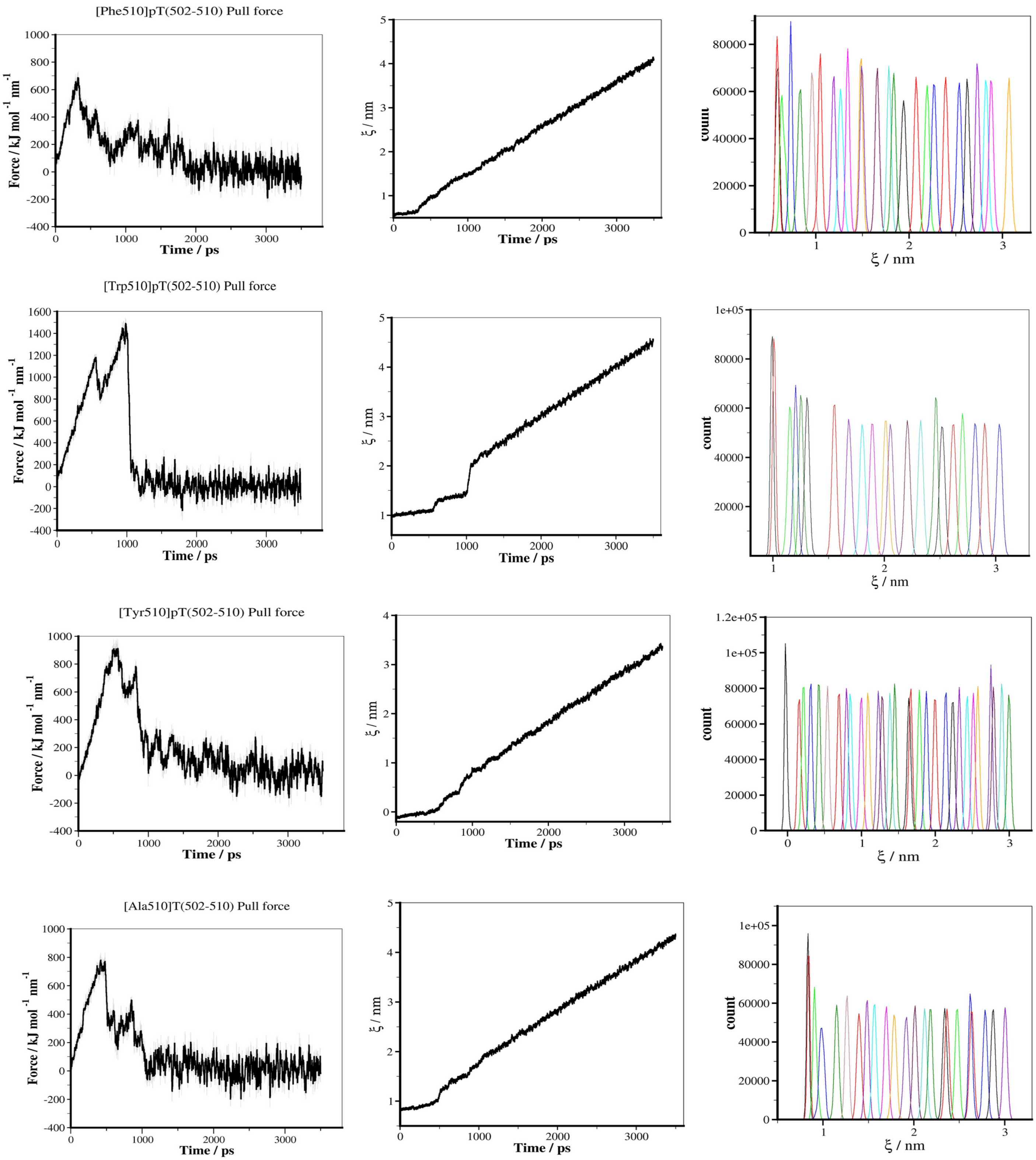
[Phe510]pT(502-510) Pull force
Force / kJ mol-1 nm-1
Time / ps
ξ / nm
Time / ps
count
ξ / nm
[Trp510]pT(502-510) Pull force
Force / kJ mol-1 nm-1
Time / ps
ξ / nm
Time / ps
count
ξ / nm
[Tyr510]pT(502-510) Pull force
Force / kJ mol-1 nm-1
Time / ps
ξ / nm
Time / ps
count
ξ / nm
[Ala510]T(502-510) Pull force
Force / kJ mol-1 nm-1
Time / ps
ξ / nm
Time / ps
count
ξ / nm

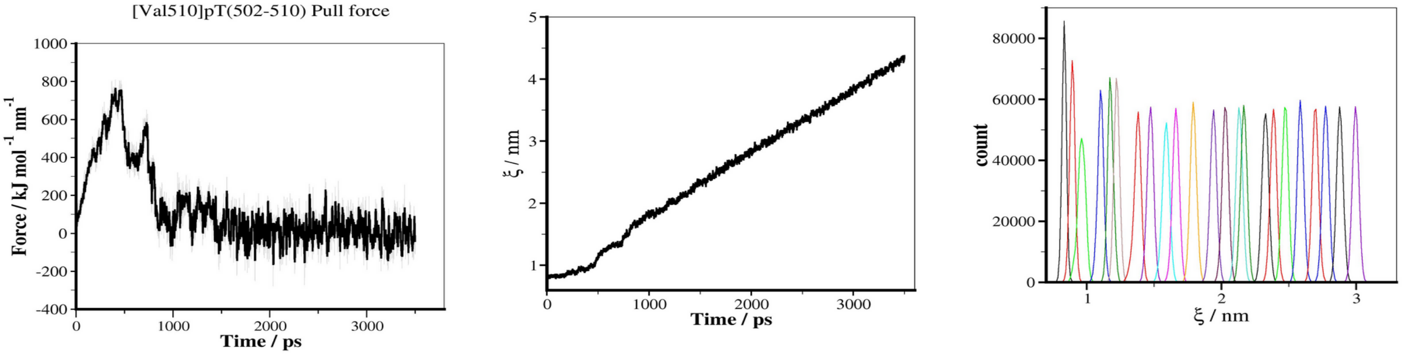


**Figure S5**. Steered molecular dynamics simulation of 14-3-3ε – peptide complexes. Time dependent external force applied to pull the peptide away from binding groove of 14-3-3ε (left panel); time dependent distance between center of mass (COM) of 14-3-3ε and COM of peptide (middle panel); umbrella histograms of configurations, each derived from 30 ns simulation (right panel).

A

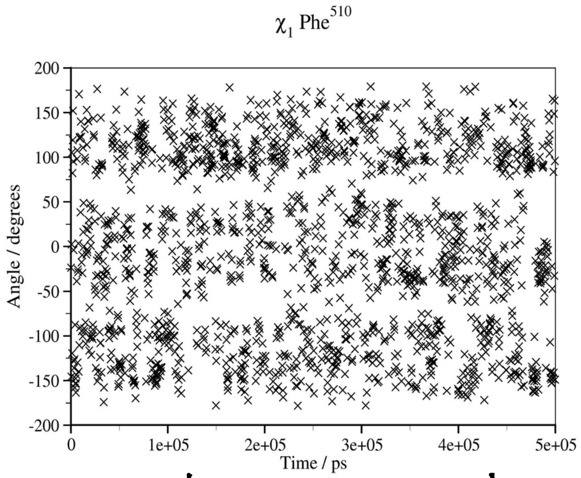


B Phe

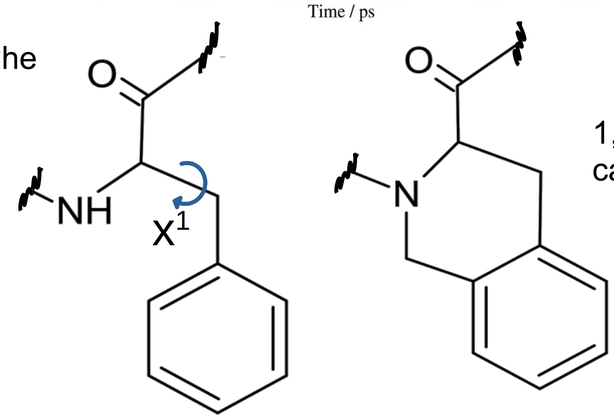


1,2,3,4 tetrahydroisoquinoline-3-carboxylic acid (Tic)

**Figure S6**. (A**) Change in** x[1] torsional angle of Phe[510] of [Phe[510]]pT(502-510) peptide during 500 ns MD simualtion. The x[1] torsional angle is flexible, the aromatic side chain fluctuates between t, *g*(+), and *g*(-) positions throughout the simulation. (B) Comparison of the structure of Phe and Tic amino acid residues, x[1] angle is indicated by a curved arrow.

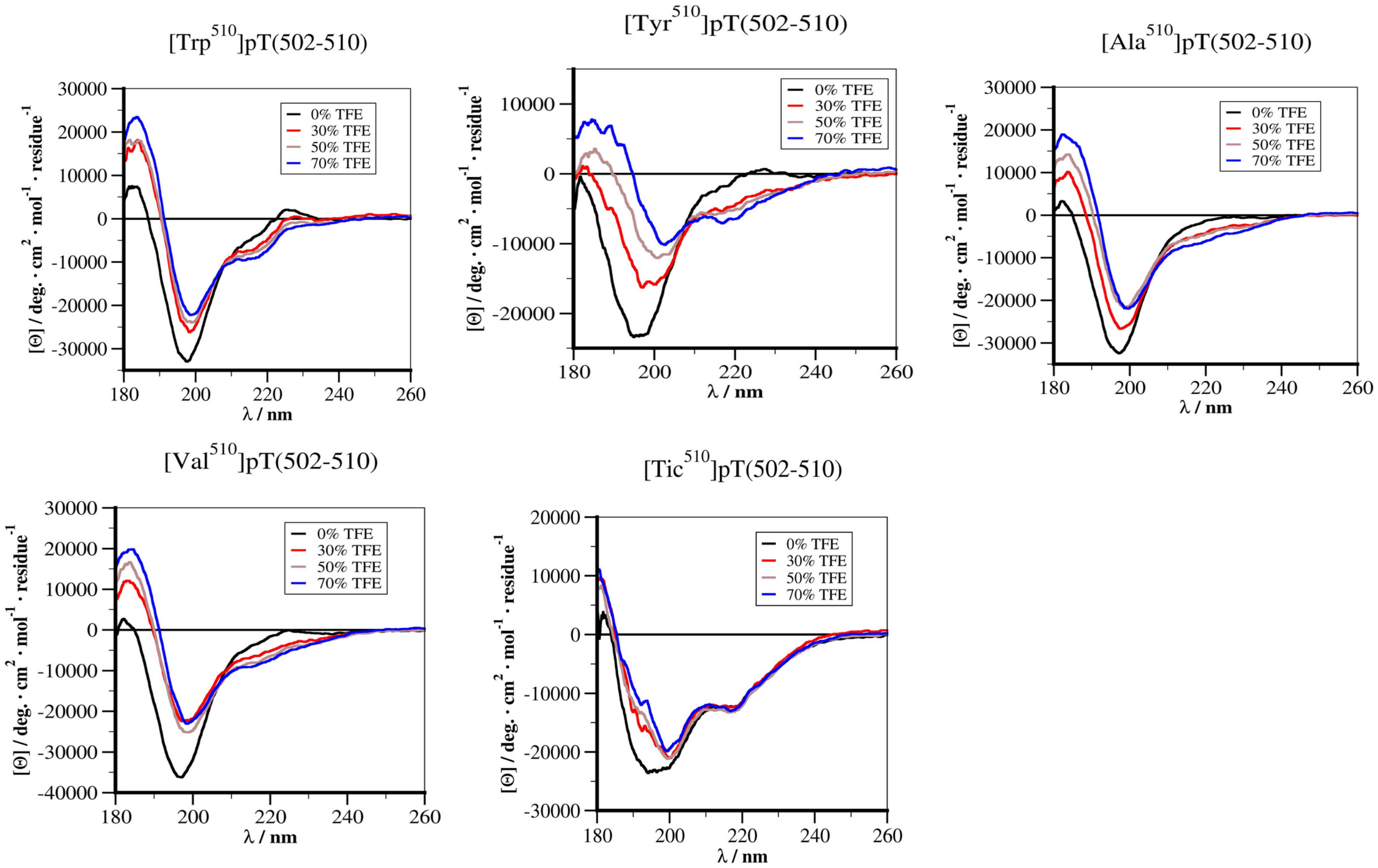


**Figure S7. E**CD spectra of peptides in different solutions. Each spectrum is an average of 20 scans. Mean residue ellipticity was determined from concentration of the peptides.

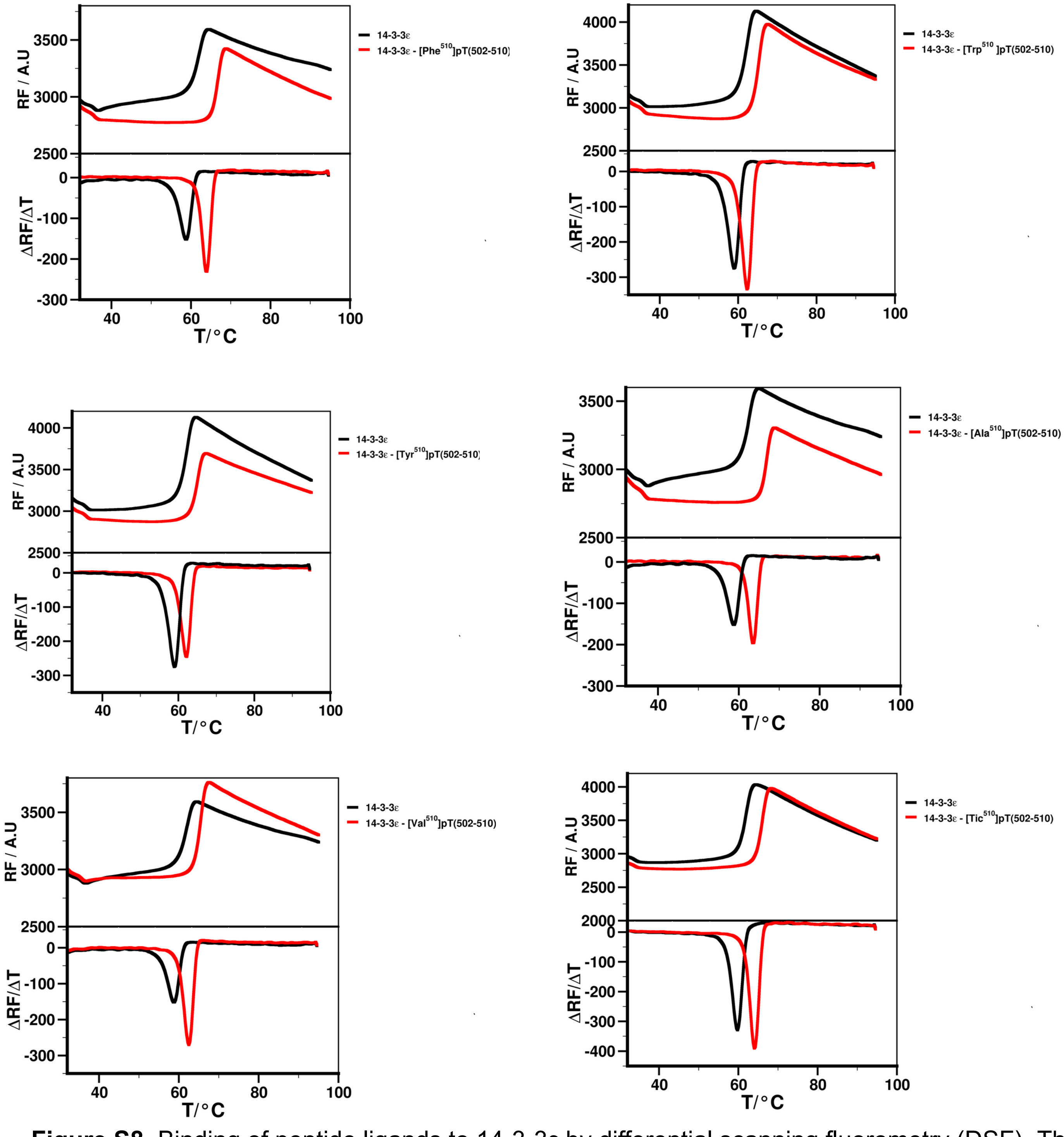


**Figure S8.** Binding of peptide ligands to 14-3-3ε by differential scanning fluorometry (DSF)**.** Thermal denaturation of 14-3-3ε without (black) and with peptide (red).

[Phe510]pT(502-510)

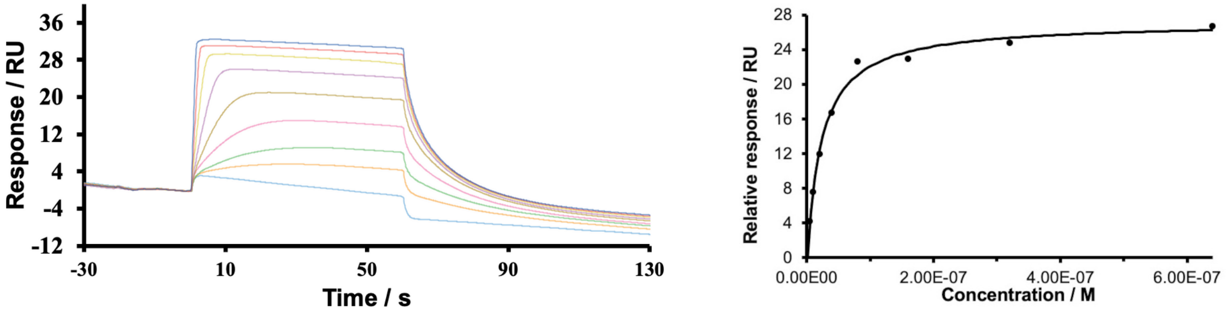


[Trp510]pT(502-510)

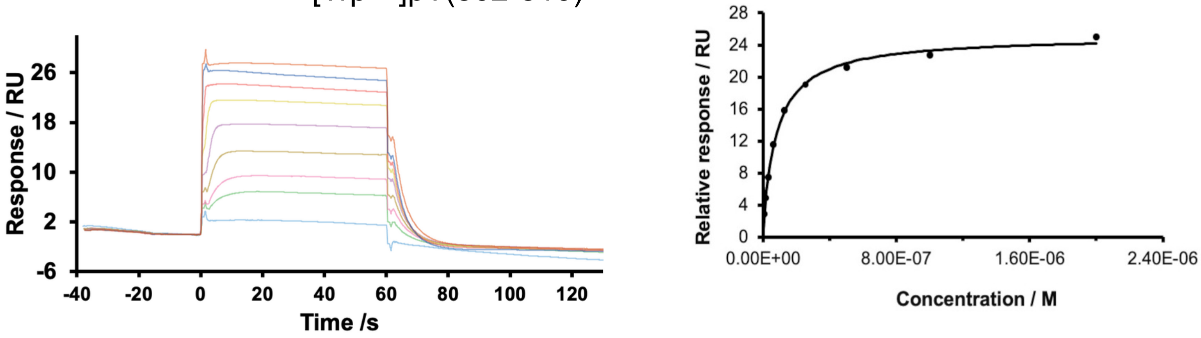


[Tyr510]pT(502-510)

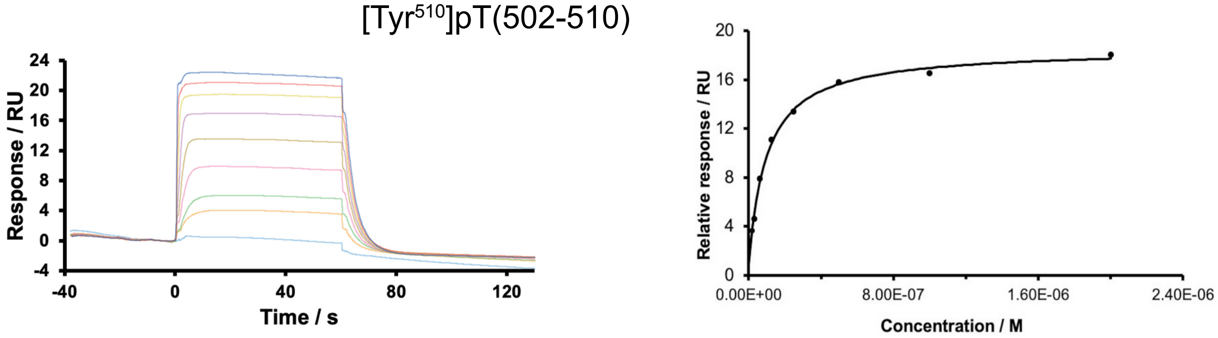


[Ala510]pT(502-510)

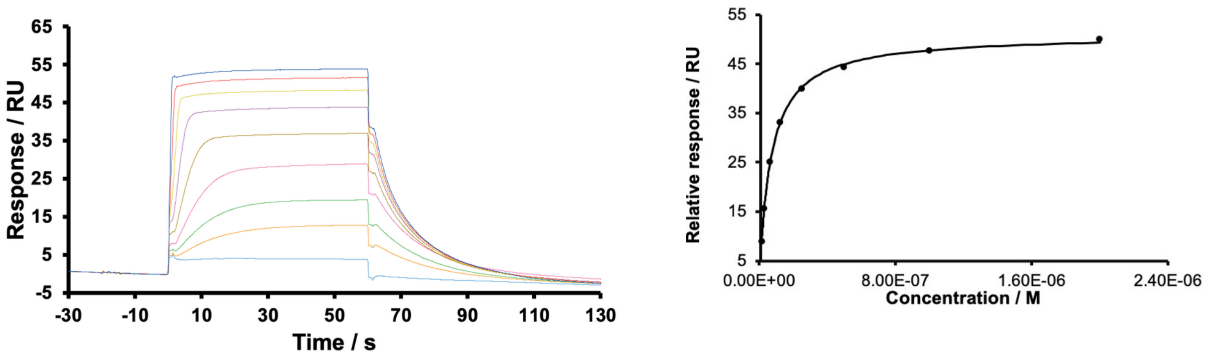

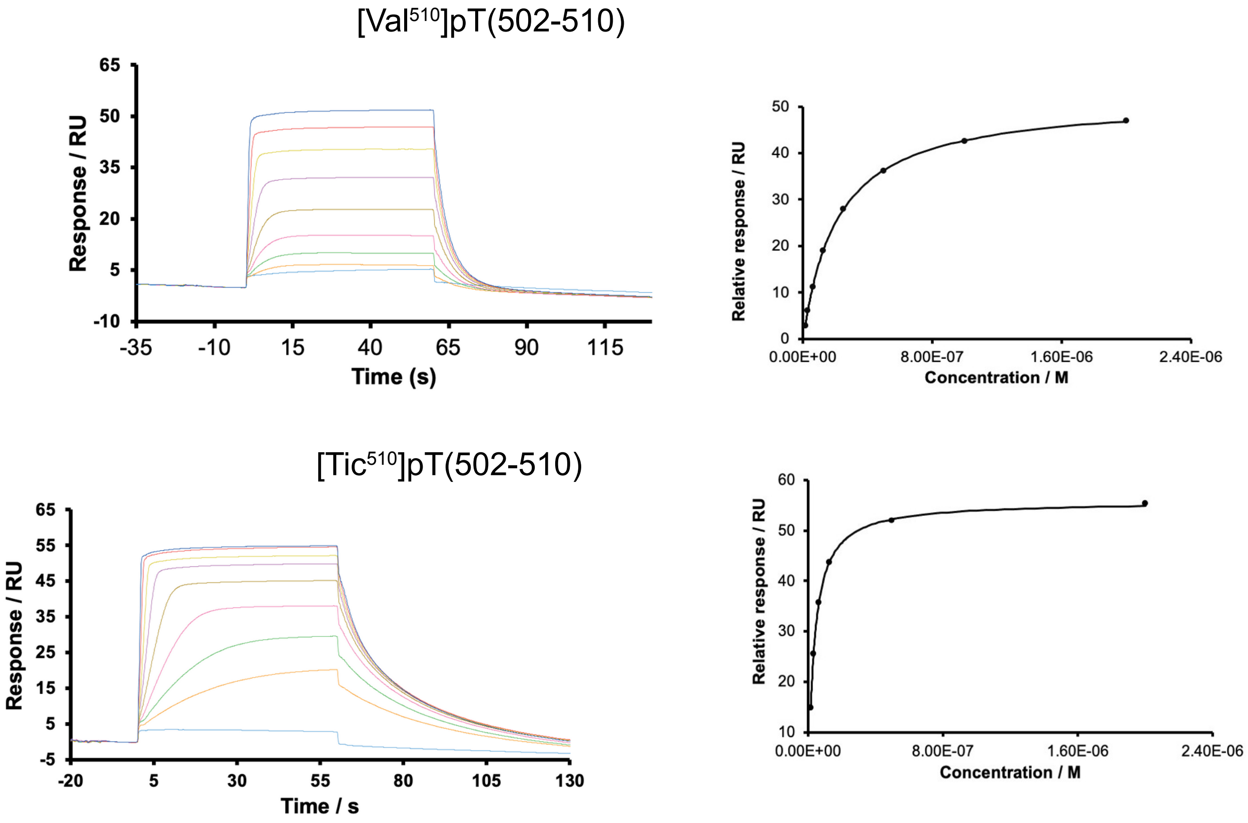


**Figure S9.** Binding of synthetic peptide ligands to 14-3-3ε by surface plasmon resonance (SPR)**.** 100 µM of 14-3-3ε was immobilized on sensor chip and the binding of peptides at different concentrations (1-2000 nM) were measured at 298 K. Sensograms and their respective binding isotherms are shown.

**Table S1. Amino acid sequence of sequentially truncated analogs of the pT peptide.**

| Peptide | Sequences* |
|---|---|
| pT | Ac-RTKSRpTWAGEKSKR-$NH_2$ |
| pT(502-514) | Ac-RTKSRpTWAGEKSK-$NH_2$ |
| pT(502-513) | Ac-RTKSRpTWAGEKS-$NH_2$ |
| pT(502-512) | Ac-RTKSRpTWAGEK-$NH_2$ |
| pT(502-511) | Ac-RTKSRpTWAGE-$NH_2$ |
| pT(502-510) | Ac-RTKSRpTWAG-$NH_2$ |
| pT(503-515) | Ac-TKSRpTWAGEKSKR-$NH_2$ |

*Ac and -$NH_2$, acetyl and amide, respectively, protecting groups.

**Table S2**. **Change in free energies of binding ($\Delta\Delta G_b$) of 14-3-3ε – peptide up on introducing modification in the pT(502-510) peptide.**

| Peptide | Sequence* | $\Delta\Delta G_b$ |
|---|---|---|
| pT(502-510) | Ac-RTKSRpTWAG-$NH_2$ | N.A |
| [$Arg^{502}$Phe]pT(502-510) | Ac-FTKSRpTWAG-$NH_2$ | −0.18 ± 0.53 |
| [$Arg^{502}$Trp]pT(502-510) | Ac-WTKSRpTWAG-$NH_2$ | 0.50 ± 0.73 |
| [$Arg^{502}$Tyr]pT(502-510) | Ac-YTKSRpTWAG-$NH_2$ | 0.67 ± 0.74 |
| [$Thr^{503}$Ala]pT(502-510) | Ac-RAKSRpTWAG-$NH_2$ | 1.88 ± 1.79 |
| [$Thr^{503}$Arg]pT(502-510) | Ac-RRKSRpTWAG-$NH_2$ | 2.44 ± 3.34 |
| [$Thr^{503}$Lys]pT(502-510) | Ac-RKKSRpTWAG-$NH_2$ | 1.85 ± 2.40 |
| [$Thr^{503}$Phe]pT(502-510) | Ac-RFKSRpTWAG-$NH_2$ | 0.93 ± 1.26 |
| [$Thr^{503}$Trp]pT(502-510) | Ac-RWKSRpTWAG-$NH_2$ | 4.80 ± 5.62 |
| [$Thr^{503}$Tyr]pT(502-510) | Ac-RYKSRpTWAG-$NH_2$ | 1.09 ± 1.98 |
| [$Ser^{505}$Ala]pT(502-510) | Ac-RTKARpTWAG-$NH_2$ | −1.00 ± 0.92 |
| [$Ser^{505}$Arg]pT(502-510) | Ac-RTKRRpTWAG-$NH_2$ | −0.15 ± 1.68 |
| [$Ser^{505}$Lys]pT(502-510) | Ac-RTKKRpTWAG-$NH_2$ | −1.23 ± 1.82 |
| [$Ser^{505}$Phe]pT(502-510) | Ac-RTKFRpTWAG-$NH_2$ | 6.25 ± 4.24 |
| [$Ser^{505}$Trp]pT(502-510) | Ac-RTKWRpTWAG-$NH_2$ | 7.06 ± 5.09 |
| [$Trp^{508}$Arg]pT(502-510) | Ac-RTKSRpTRAG-$NH_2$ | 9.63 ± 2.20 |
| [$Trp^{508}$Lys]pT(502-510) | Ac-RTKSRpTKAG-$NH_2$ | 7.495 ± 1.855 |

| | | |
|---|---|---|
| [Trp$^{508}$Phe]pT(502-510) | Ac-RTKSRpTFAG-$NH_2$ | −0.31 ± 2.66 |
| [Trp$^{508}$Tyr]pT(502-510) | Ac-RTKSRpTKAG-$NH_2$ | 0.94 ± 2.30 |
| [Ala$^{509}$Arg]pT(502-510) | Ac-RTKSRpTWRG-$NH_2$ | −1.72 ± 0.90 |
| [Ala$^{509}$Lys]pT(502-510) | Ac-RTKSRpTWKG-$NH_2$ | −1.74 ± 0.57 |
| [Ala$^{509}$Phe]pT(502-510) | Ac-RTKSRpTWFG-$NH_2$ | −2.25 ± 0.88 |
| [Ala$^{509}$Pro]pT(502-510) | Ac-RTKSRpTWPG-$NH_2$ | −1.79 ± 0.74 |
| [Ala$^{509}$Trp]pT(502-510) | Ac-RTKSRpTWWG-$NH_2$ | −1.95 ± 0.65 |
| [Ala$^{509}$Tyr]pT(502-510) | Ac-RTKSRpTWKG-$NH_2$ | −2.05 ± 0.72 |

Modified residues are highlighted in red. The $\Delta\Delta G_b$ was calculate by FoldX program, the error was evaluated over n=5 independent calculations.
*Ac and -$NH_2$, acetyl and amide, respectively, protecting groups.

**Table S3. Composition of fraction of secondary structure content of peptides.**

| Peptides | Helix | | β-strand | | β-turn | | unordered | | total | |
|---|---|---|---|---|---|---|---|---|---|---|
| | $H_2O$ | 70% TFE | $H_2O$ | 70% TFE | $H_2O$ | 70% TFE | $H_2O$ | 70% TFE | $H_2O$ | 70% TFE |
| pT | 0 | 0.36 | 0.1 | 0.19 | 0.05 | 0.12 | 0.83 | 0.34 | 0.97 | 1.0 |
| pT(502-510) | 0 | 0.02 | 0.13 | 0.09 | 0.08 | 0.04 | 0.77 | 0.83 | 0.98 | 0.99 |
| [Phe$^{510}$]pT(502-510) | 0.02 | 0.06 | 0.24 | 0.33 | 0.16 | 0.16 | 0.56 | 0.45 | 0.98 | 1 |
| [Trp$^{510}$]pT(502-510) | -0.02 | 0.02 | 0.19 | 0.29 | 0.09 | 0.12 | 0.71 | 0.56 | 0.97 | 0.99 |
| [Tyr$^{510}$]pT(502-510) | -0.02 | 0.10 | 0.13 | 0.18 | 0.08 | 0.13 | 0.78 | 0.59 | 0.97 | 1 |
| [Ala$^{510}$]pT(502-510) | -0.02 | 0.06 | 0.06 | 0.15 | 0.03 | 0.08 | 0.91 | 0.69 | 0.98 | 0.99 |
| [Val$^{510}$]pT(502-510) | -0.02 | 0.06 | 0.08 | 0.19 | 0.04 | 0.13 | 0.88 | 0.62 | 0.98 | 1 |
| Tic$^{510}$]pT(502-510) | 0.07 | 0.18 | 0.39 | 0.34 | 0.21 | 0.22 | 0.32 | 0.27 | 0.99 | 1.01 |

Secondary structure of peptides as determined by CD spectropolarimetry in $H_2O$ and in 70 % TFE. Components of secondary structures of peptides were obtained by deconvoluting CD spectra using DSSTR method of DichroWeb CD analysis web server.

**Table S4**. **Kinetics of the peptide binding to 14-3-3ε.**

| **Peptide** | **$k_{on}$ / $M^{-1}$ $s^{-1}$ (*$10^6$)** | **$k_{off}$ / $s^{-1}$ (*$10^{-1}$)** | **$K_D$ / nM** |
|---|---|---|---|
| pT | 0.84 ± 0.075 | 1.26 ± 0.06 | 149.92 ± 6.58 |
| pT(502-510) | 1.55 ± 0.35 | 1.89 ± 0.89 | 118.89 ± 31.52 |
| [$Phe^{510}$]pT(502-510) | 2.21 ± 0.92 | 0.45 ± 0.06 | 20.78 ± 2.73 |
| [$Trp^{510}$]pT(502-510) | 2.38 ± 0.28 | 2.015 ± 0.16 | 85.41 ± 10.94 |
| [$Tyr^{510}$]pT(502-510) | 2.49 ± 0.43 | 2.08 ± 0.43 | 83.64 ± 5.67 |
| [$Ala^{510}$]pT(502-510) | 1.01 ± 0.14 | 0.84 ± 0.01 | 84.22 ± 13.04 |
| [$Val^{510}$]pT(502-510) | 0.83 ± 0.048 | 1.96 ± 0.21 | 229.74 ± 13.15 |
| [$Tic^{510}$]pT(502-510) | 1.29 ± 0.17 | 0.43 ± 0.03 | 33.70 ± 2.55 |

The $k_{on}$ and $k_{off}$ were obtained by fitting the sensograms to1:1 binding model, $K_D$ was calculated as the ratio of $k_{off}$ and $k_{on}$. The values correspond to the average and SD of n ≥ 3.

flurot
